# Nuclear NFκB Activity Balances Purine Metabolism in Cellular Senescence

**DOI:** 10.1101/2023.04.18.536673

**Authors:** Sho Tabata, Keita Matsuda, Kenshiro Nagai, Yoshihiro Izumi, Masatomo Takahashi, Yasutaka Motomura, Ayaka Ichikawa Nagasato, Shuichi Shimma, Kazuyo Moro, Takeshi Bamba, Mariko Okada

## Abstract

Upregulation of nuclear factor κB (NFκB) signaling is a hallmark of aging and major cause of age-related chronic inflammation; however, its physiological functions and mechanisms remain unclear. By combining mathematical modeling and experiments, we show that dysfunction of negative feedback regulators of NFκB, IκBα and A20, alters the NFκB nuclear dynamics from oscillatory to sustained, promoting cellular senescence by remodeling epigenetic regulation and metabolic landscape. Sustained NFκB activity by IκBα downregulation enhanced inflammation- and senescence-associated gene expression through increased NFκB-DNA binding and slowed the cell cycle by upregulating purine catabolism via mTORC2/AKT pathways. Notably, IκBα knockdown combined with A20 overexpression resulted in lower NFκB amplitude, cytokine expression, and SA-β-gal activity than IκBα knockdown alone. IκBα downregulation is correlated with hypoxanthine phosphoribosyltransferase 1 (HPRT1) expression in the purine salvage pathway in aged mouse hearts. Our study suggests that nuclear NFκB homeostasis is critical for balancing purine metabolism associated with chronic inflammation and tissue aging.

## INTRODUCTION

Aging is a complex and irreversible phenomenon that reflects lifelong biochemical responses to the environment. Recent research has shown that age-related dysfunction occurs from the genome to organ levels ^1, 2^. Inflammatory aging, which refers to aging-associated chronic inflammation, critically impacts the induction of a wide range of age-related human diseases ^3^, and its regulation is becoming increasingly important in aging societies. Elderly individuals exhibit low-grade and persistent chronic inflammation in their tissues along with elevated levels of circulating inflammatory factors that activate nuclear factor κB (NFκB) signaling, such as interleukin-6 and tumor necrosis factor a (TNFα) ^4, 5^. Meanwhile, the accumulation of senescent cells during aging produces various pro-inflammatory cytokines that represent a senescence-associated secretory phenotype (SASP), which increases the risk of developing age-related diseases ^6^. As NFκB is the master transcription factor for SASP genes ^7, 8^, NFκB is considered a positive feedback regulator in inflammatory aging and cellular senescence. However, several studies have also shown that NFκB prevents cellular senescence, therefore it remains unclear which properties of NFκB activity is related to cellular senescence ^9, 10^. Herein, we speculate that a quantitative rather than qualitative understanding of NFκB activity is necessary to unravel the role of NFκB in cellular senescence and inflammatory aging.

NFκB regulates the expression of genes involved in immune responses and cell proliferation, apoptosis, and senescence ^11–13^. Excessive NFκB activity induces pathological inflammation and contributes to the progression of various human diseases ^14^. Notably, NFκB also induces transient, sustained, or oscillatory nuclear dynamics associated with different biological functions and cell fates ^15^. The quantitative features of NFκB activity, including the peak amplitude, duration, speed, and oscillation, are associated with the discrimination of immune threats by macrophages ^16^, and it encodes a short-term history derived from prior pathogenic and cytokine signals, affecting later cell functions ^17^. These studies demonstrate that the homeostasis of NFκB nuclear dynamics is important for the maintenance of cellular functions.

The canonical NFκB pathway undergoes negative feedback regulation in response to environmental factors and exhibits stimulus-specific oscillatory dynamics. In particular, NFκB inhibitor alpha (IκBα), which is rapidly expressed upon NFκB activation, acts as a strong negative feedback regulator by binding to NFκB and inhibiting its nuclear translocation, thereby inducing characteristic NFκB oscillatory patterns ^18^. Notably, knockout (KO) or knockdown (KD) of the IκBα gene in TNFα-stimulated cells results in sustained NFκB nuclear activation and upregulation of immediately expressed genes that show distinct patterns generated from oscillatory NFκB ^19–21^. However, the physiological significance of NFκB dynamics and their long-term implications on signaling crosstalk, epigenetics, and metabolic regulation have not been fully addressed. In this study, we investigated the impact of changes in TNFα-induced NFκB dynamics on cell functions and found that sustained NFκB induces activation of purine catabolic pathways *via* the mTORC2/AKT pathway and prolongs the cell cycle period, rewires epigenetic states by enhanced NFκB-DNA binding to promote inflammatory gene expression. These results indicate that sustained nuclear NFκB activity significantly alters the landscape of the cellular network to induce cellular senescence and inflammatory aging.

## RESULTS

### Sustained NFκB activity induces growth arrest and cellular senescence

First, we confirmed that IκBα KD alters oscillatory NFκB nuclear activity to sustain activity in MCF-7 cells stimulated with TNFα (Figures 1A, S1A, and S1B). The change in NFκB dynamics suppressed cell proliferation after 24 hr of TNFα stimulation (Figure 1B) and led to a flattened cell shape and significantly increased cell size (Figures 1C, S1C, and S1D). These features are similar to the senescence phenotype ^22^. Therefore, we examined SA-β-gal activity, one of the indicators of cellular senescence ^23^. Although TNFα alone did not promote SA-β-gal activity, under IκBα KD/KO conditions, TNFα increased the number of SA-β-gal positive cells after 24 hr (Figures 1D and S1E–S1G). The number of SA-β-gal positive cells was suppressed by KD of RELA, a subunit of NFκB, in IκBα KD+TNFα conditions (Figures 1E and S1H), thus confirming that the phenomenon is NFκB-dependent. As senescent cells typically exhibit a SASP ^6, 23^, we next examined the effects under IκBα KD+TNFα conditions. The expression of various SASP genes, including *IL8* and *CCL2*, was drastically elevated (Figures 1F and 1G). The amount of Cyclin D1, which is essential for the G1/S transition, was decreased compared to that with TNFα alone (Figure 1H). Furthermore, a time-course cell cycle analysis using FUCCI live cell imaging revealed that IκBα KD+TNFα increases the number of the cells at the G2/M phase after 24 hr and prolongs the duration of the G2/M phase (median values: control, 3.8; IκBα KD, 8.3; *P* < 0.05) (Figures 1I, S1I, and S1J). These results indicate that the sustained NFκB activity mimicked by IκBα KD+TNFα modulates cell cycle checkpoints and inhibits cell proliferation.

**Figure 1.**
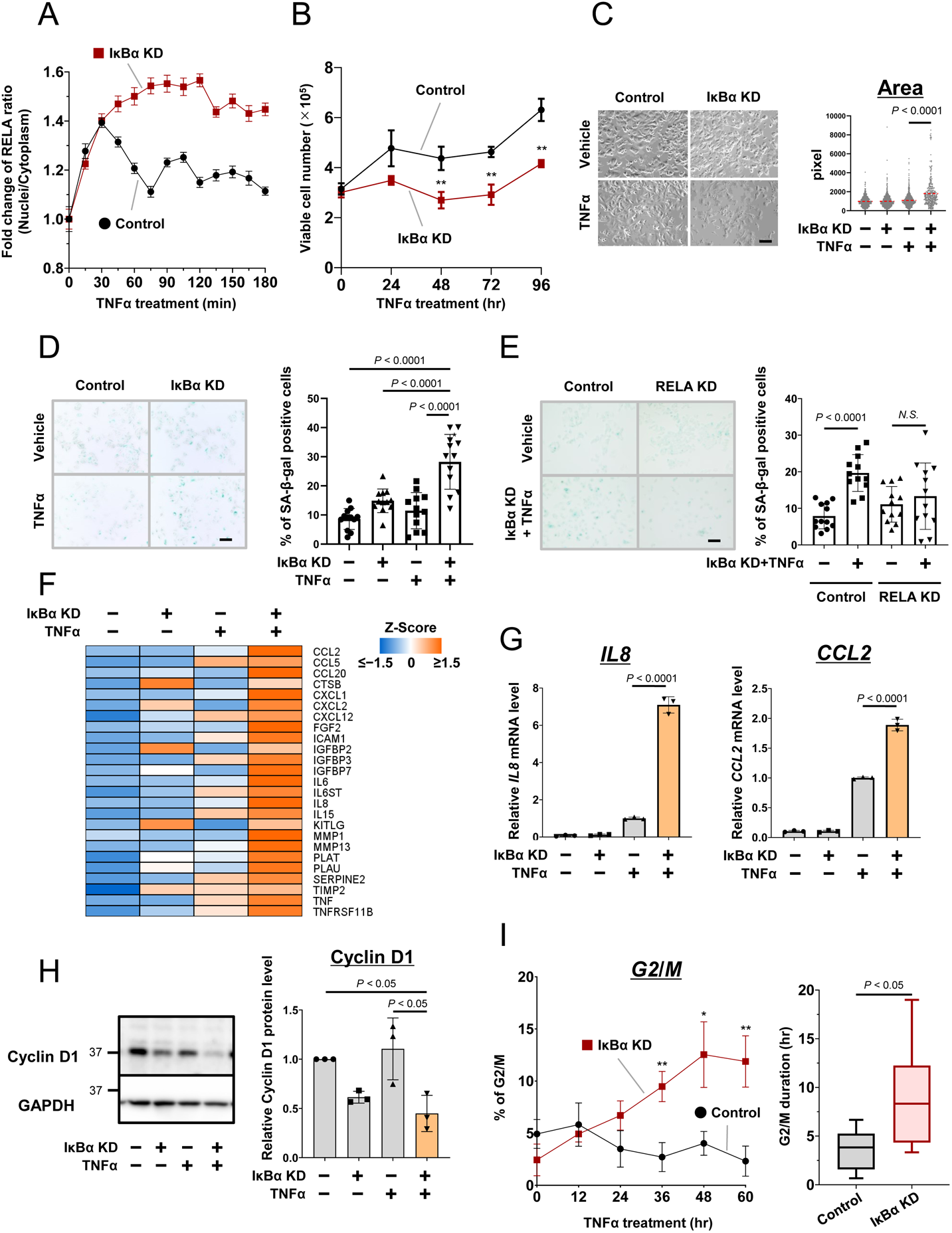
Effect of NFκB inhibitor alpha (IκBα) knockdown on growth arrest and cellular senescence in MCF-7 cells treated with tumor necrosis factor alpha (TNFα). (A) Time-course of nuclear factor κB (NFκB) (RELA) abundance in the control and IκBα knockdown (KD) MCF-7 cells treated with TNFα (10 ng/mL) (see the Methods). (B) Effect of IκBα KD on cell viability. (C) Changes in cell morphology in the IκBα KD MCF7 cells treated with TNFα for 24 hr. Scale bars, 100 μm. (D) Effect of IκBα KD on SA-β-gal staining in MCF-7 cells treated with TNFα for 24 hr. Scale bars, 100 μm. (E) Effect of RELA KD on the IκBα KD-induced SA-β-gal staining. Scale bars, 100 μm. (F and G) Senescence-associated secretory phenotype (SASP) gene expression in MCF-7 cells under IκBα KD+TNFα conditions. Cells were treated with TNFα for 24 hr. The mRNA level of SASP genes was measured by RNA-seq (F) and qPCR (G). (H) Effect of IκBα KD on the protein expression of Cyclin D1 in MCF-7 cells treated with TNFα for 48 hr. (I) G2/M ratio (left) and duration (right) in TNFα-treated control and IκBα KD MCF7 cells. Data represent the mean ± SD [(B), (D), (E), (G), (H) and (I, left)] or ± SE (A), median and interquartile range (25th and 75th percentiles), with error bars showing the minimum to maximum (I, right). *P* values were determined using unpaired two-tailed Student’s t tests [(B) and (I)] or one-way ANOVA followed by Tukey’s test [(C), (D), (E), (G), and (H)]. \**P* < 0.05, \*\**P* < 0.01. *N*.*S*., not significant.

Of note, a previous study showed that IκBα not only plays a negative role in NFκB activity but also positively affects the induction of several NFκB target genes ^19^. Therefore, we sought to determine whether the sustained NFκB dynamics were responsible for the induction of cellular senescence rather than IκBα itself. In addition to IκBα, A20 has been shown to act as a negative feedback regulator of the NFκB pathway by inhibiting upstream IKK activity and TTR complex (TRAF2, TRADD and RIP) binding to the TNFα receptor and modulating the oscillatory dynamics of NFκB ^24^. Although the regulatory mechanisms of IκBα and A20 on the NFκB pathway are different, downregulation of each molecule leads to sustained NFκB dynamics ^24, 25^. Therefore, we investigated whether A20 KD promotes cellular senescence. A20 KD+TNFα indeed exhibited sustained NFκB activity (Figures 2A and S2A), increased SA-β-gal staining and SASP gene expression (*IL8* and *CCL2*) (Figures 2B and 2C), and slightly reduced Cyclin D1 expression (Figure 2D) in MCF-7 cells; however, these effects were weaker than those of IκBα KD (Figures 1A and 1D–1H).

**Figure 2.**
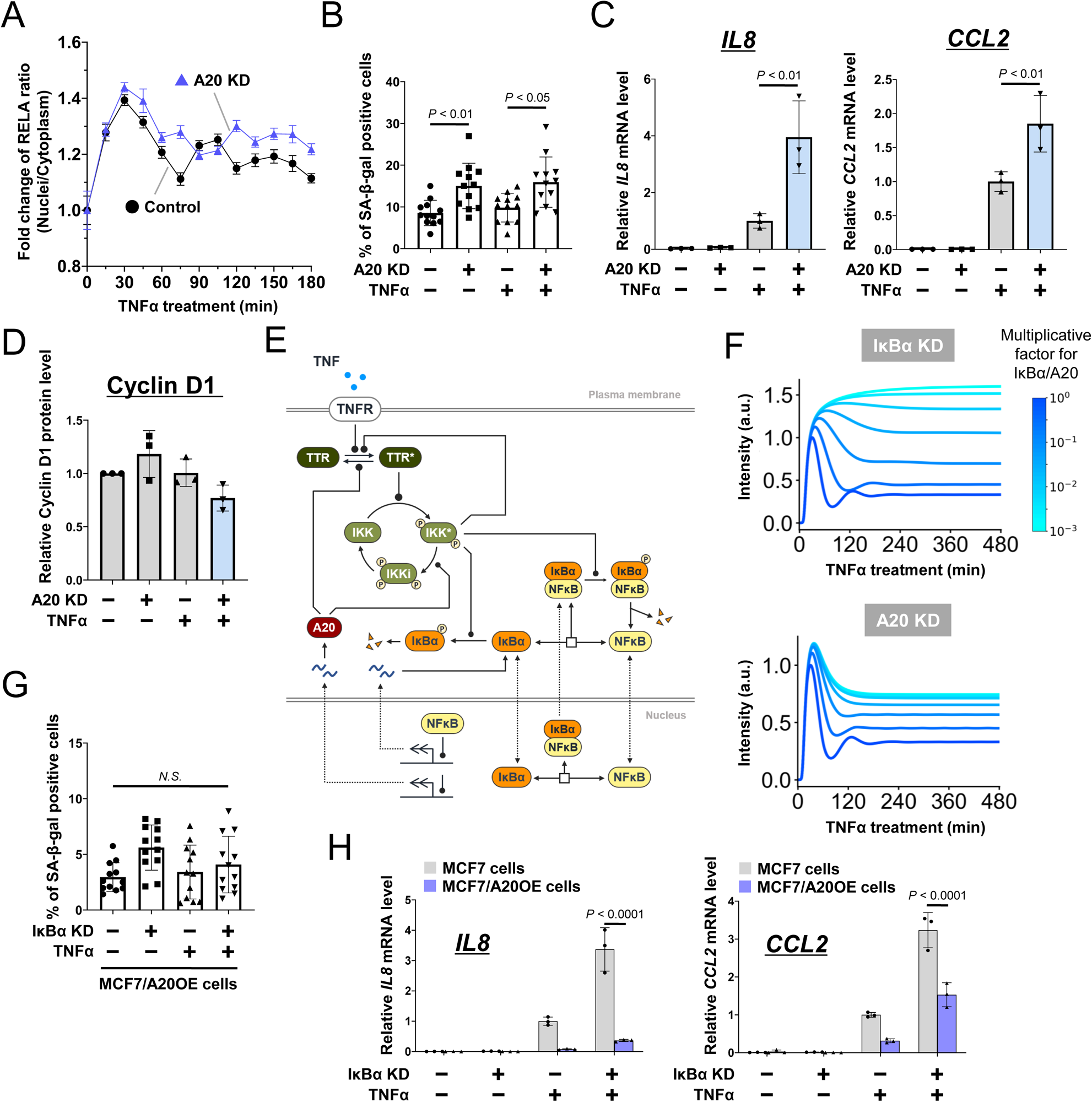
Mathematical modeling of NFκB dynamics and effects of IκBα and A20 knockdown. (A) Time-course nuclear NFκB (RELA) abundance in A20 knockdown (KD) MCF-7 cells treated with TNFα for the indicated time. The control is the same dataset as in Figure 1A. (B–D) Effect of A20 KD on SA-β-gal staining (B), SASP gene expression (C), and Cyclin D1 protein expression (D) in MCF-7 cells treated with TNFα (10 ng/mL). (E) Diagram of the TNFα-NFκB signaling network. (F) Simulations of nuclear NFκB abundance in IκBα (upper) and A20 KD (lower) cells treated with TNFα for the indicated time. The color bar indicates the fold change values in IκBα or A20 transcription rate compared to that in the control. (G and H) Effect of IκBα KD in SA-β-gal staining (G) and SASP gene expression (H) in A20-overexpressing MCF-7 cells treated with TNFα. Data represent the mean ± SD [(B), (C), (D), (G) and (H)] or ± SE (A). *P* values were determined using the one-way ANOVA followed by Tukey’s test [(B), (C), (G), and (H)]. *N*.*S*., not significant.

To evaluate the quantitative difference in the effects of IκBα and A20 on NFκB nuclear dynamics, we constructed a mathematical model of the NFκB pathway (Figures 2E, S2B and Tables S1–3) modified from earlier studies ^26, 27^ and performed simulation analyses. We trained the model with time-course western blot data obtained from TNFα-stimulated MCF-7 cells (Figures S2C–S2F) and analyzed the NFκB dynamics altered by IκBα and A20 KD. The simulated downregulation or overexpression (OE) of IκBα or A20 showed that IκBα KD has a greater influence on changing nuclear NFκB activity from oscillatory to sustained, thereby inducing a higher total amount of nuclear NFκB activity (Figures 2F and S2G). Sensitivity analysis (see the Methods) also confirmed that IκBα-related reactions presented slightly higher sensitivity to the sum of nuclear NFκB activity than the A20 reactions (Figure S2H), suggesting that IκBα has a more critical effect on nuclear localization of NFκB. The result was consistent with the experimental results of NFκB nuclear dynamics (Figures 1A and 2A) and the extent of cellular senescent phenotypes (Figures 1D–1H and 2B–D) between IκBα and A20 KD. These results indicate that the total nuclear abundance of NFκB associated with prolonged NFκB activity may be critical in the induction of cellular senescence. To further evaluate the importance of these quantitative features of NFκB (e.g., oscillation or abundance) and determine whether A20 OE could compensate for the effect of IκBα downregulation to recover oscillation dynamics to inhibit cell senescence, we first performed simulation analyses of nuclear NFκB dynamics by combining IκBα KD and A20 OE (Figure S2I). Under A20 OE, IκBα KD still exhibited sustained NFκB activity to some extent; however, its overall amplitude was reduced. To confirm this result experimentally, we established A20 OE MCF-7 cells (Figure S2J) and examined the effect of IκBα KD on NFκB dynamics. As computationally predicted, A20 OE reduced the amplitude of NFκB activity in IκBα KD+TNFα conditions but failed to fully alter the NFκB dynamics from sustained to oscillatory (Figure S2K). However, under the same conditions, the increase in the number of SA-β-gal positive cells associated with IκBα KD (Figure 1D) was mitigated by A20 OE (Figures 2G), which also reduced *IL8* and *CCL2* expression (Figure 2H). These results suggest that the amount of nuclear NFκB level is more critical for the induction of cellular senescence.

### Transcriptional regulation of sustained NFκB activity *in vitro* and *in vivo*

Subsequently, we investigated the relationship between NFκB nuclear dynamics and target gene regulation. First, we extracted differentially expressed genes (DEGs) in MCF7 cells between IκBα KD+TNFα (48 hr) and TNFα conditions and identified 687 upregulated DEGs, including cellular senescence-promoting and SASP genes (e.g., *CD36*, *BCL3*, *CCL2*, and *S100A8/A9*) ^1, 23, 28, 29^ (Figure 3A). These DEGs were classified into four clusters based on the time-course patterns of gene expression, which was measured every 15 min under the IκBα KD+TNFα condition ^19^ (Figure 3B). To clarify whether the gene expression time-course was associated with the chromatin status, chromatin accessibility was assessed using an assay for transposase-accessible chromatin sequencing (ATAC-seq) data. We found that the ATAC signal was elevated in the flanking regions of genes in cluster 2, which showed a monotonically increasing pattern, and their signals were stronger in the IκBα KD+TNFα condition than the TNFα or IκBα KD alone condition (Figure 3C). Considering that the chromatin accessibility enhanced by IκBα KD+TNFα might be critical for the regulation of genes controlled by NFκB, we extracted the regions where chromatin accessibility was significantly enhanced by IκBα KD+TNFα (Figure S3) and assessed the NFκB binding to these regions. We analyzed NFκB (RELA)-chromatin immunoprecipitation sequencing (ChIP-seq) data and found that the RELA signal intensity in chromatin open regions proximal to cluster 2 genes was markedly increased under IκBα KD+TNFα conditions; however, it was not observed under TNFα alone (Figure 3D). Furthermore, we investigated whether the number of NFκB bindings affects gene expression in each cluster ^30^. Notably, the average number of RELA binding sites in cluster 2 genes was significantly higher than that for all NFκB target genes (Figure 3E). Since cluster 2 genes were enriched with several cytokine signaling pathways, such as TNFα and IL17 (Figure 3F), these results suggest that an inflammatory positive feedback loop is epigenetically enhanced by sustained nuclear NFκB localization.

**Figure 3.**
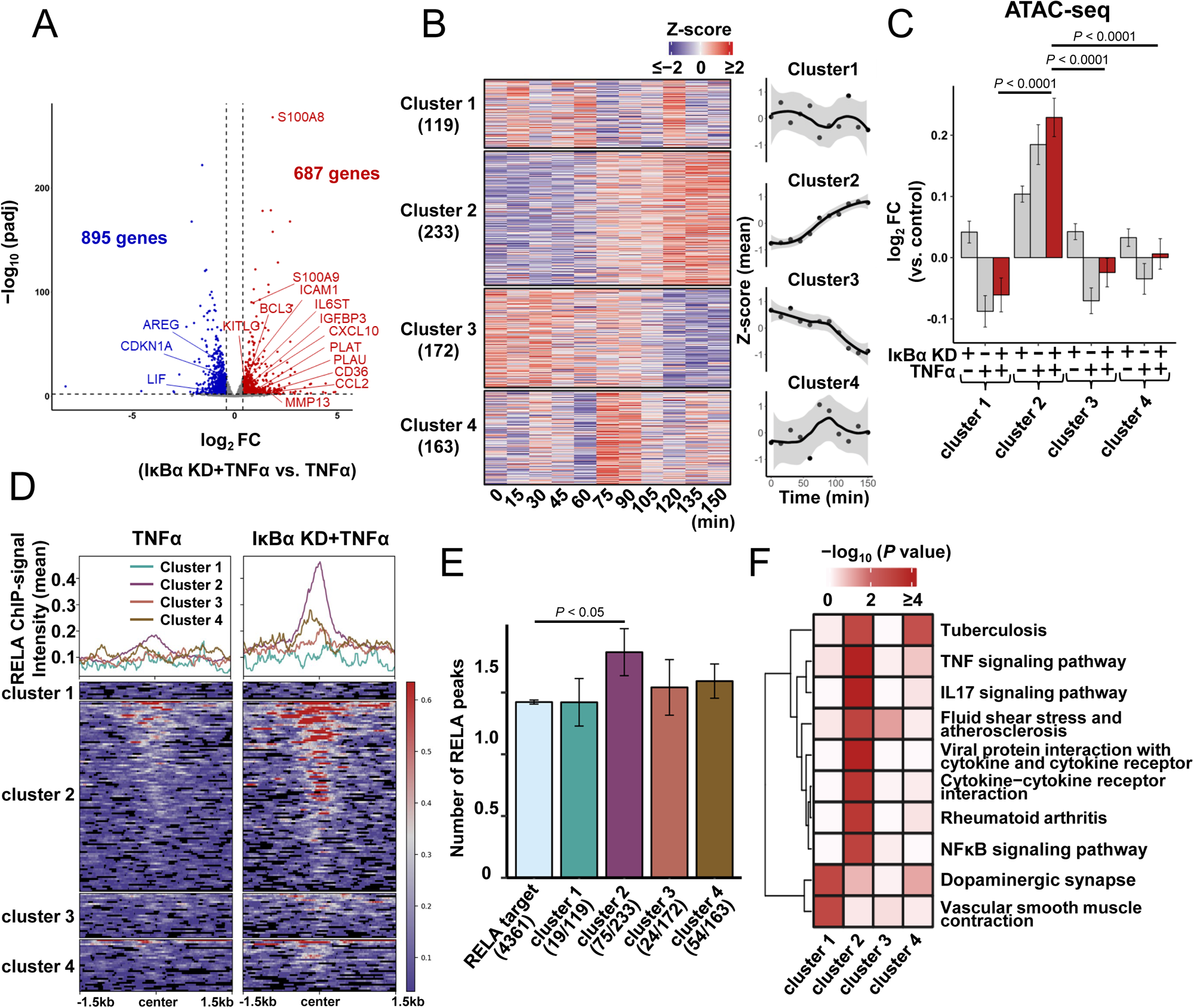
Transcriptional regulation by sustained NFκB activation. (A) Volcano plot of differentially expressed genes (DEGs) between TNFα alone and IκBα KD+ TNFα in MCF-7 cells. Cells were treated with TNFα for 48 hr. Red dots indicate DEGs identified by adjusted *P* value (padj)L<L0.01 and log2 fold change (log2 FC)L>L0.4. Blue dots indicate DEGs identified by padjL<L0.01 and log_2_ FCL<−0.4. Among the DEGs, the cellular senescence-promoting and SASP genes are labeled. (B) Time-course clustering of mRNA level (Z-scored log_2_TPM) in IκBα KD MCF-7 cells stimulated with TNFα (10 ng/mL) up to 150 min. Lines on the right indicate approximate curves, and colored areas indicate 95% confidence intervals. (C) Effect of IκBα KD on chromatin accessibility for genes in each cluster in MCF-7 cells treated with TNFα (48 hr). Data represent the log_2_ fold change (FC) vs. control conditions (D) Effect of IκBα KD and TNFα (120 min) on RELA ChIP signal intensity in open chromatin regions. Row in the heatmap indicates the signal intensity in each region. Lines in the line chart shows the average RELA signal intensity. (E) Average count of RELA ChIP peaks for NFκB target genes in each cluster. Genes with RELA peaks in the vicinity of TSS were defined as NFκB target genes (4,361 genes). NFκB target genes in each cluster were identified (cluster 1, 19 genes; cluster 2, 75 genes; cluster 3, 24 genes; cluster 4, 54 genes). (F) Heatmap of KEGG enrichment analysis of each cluster genes identified in Figure 3B. Data represent the mean ± SE [(C) and (E)]. *P* values were determined using one-way ANOVA followed by Tukey’s test [(C) and (E)].

Senescent cells accumulate in aged tissues in a variety of mammals ^23^. Given that sustained NFκB activity may be involved in promoting cellular senescence and chronic inflammation associated with aging *in vivo*, we examined the levels of IκBα, A20, and TNFα in young (8 weeks old) and aged (69–73 weeks old) mice (Figures S4A–S4D). We found decreased IκBα and A20 protein expression and increased TNFα in the heart tissues of aged mice (Figure 4A). We also found a marked increase in the NFκB-positive area in the cell nuclei of aged hearts using immunohistochemistry staining (Figures 4B and S4E) and upregulation of the cellular senescence markers *p16* and *Il-6* (Figure 4C). Since p16 is a negative regulator of CDK4/6, these data imply that cytokine signaling and cell cycle delays are increased in association with persistent NFκB activity. RNA-seq analysis of heart tissues from young and aged mice and predictions of active transcription factors using DoRothEA ^31^ suggested that NFκB family members NFKB1 and RELA were activated in aged tissue (Figure 4D). The fold change relationship between genes with sustained NFκB activity *in vitro* (IκBα KD+TNFα vs. control MCF-7) and those showing aging *in vivo* (aged vs. young mouse hearts) revealed that in MCF-7 cells, the expression of cluster 2 genes, which contain cellular senescence-promoting and SASP genes, was correlated with the expression of age-related genes in the heart (R = 0.31, *P* < 0.0001) (Figure 4E). In addition, we examined the relationship between the number of NFκB binding sites and the increase in gene expression with aging in the mouse heart. After predicting putative NFκB targets within ± 500 bp from the transcription start site (TSS) using the ChIP-Atlas ^32^ (see the Methods), we classified the genes into four groups (no, low, medium, and high upregulation) according to the degree of fold change in expression (aged vs. young mice) (Figure 4F). Counting NFκB binding sites in the flanking regions (within ± 10,000 bp from TSS) of the genes using HOMER ^33^ revealed that the “high” expression group included a significantly higher number of NFκB binding motifs (*P* < 0.0001, high vs. no upregulation) (Figure 4G). Gene Ontology (GO) analysis of the high group genes in the aged heart (Figure 4H) showed a high enrichment of inflammatory signals, which was similar to the GO results for the cluster 2 gene set in MCF-7 cells (Figure 3F). These results suggested that sustained NFκB activity might contribute to inflammatory progression in the hearts of aged mice by upregulating the expression of senescence-related genes with increased DNA-NFκB binding, and this NFκB-mediated transcriptional regulation is common between aging tissue and the senescent cells mimicked by IκBα KD+TNFα.

**Figure 4.**
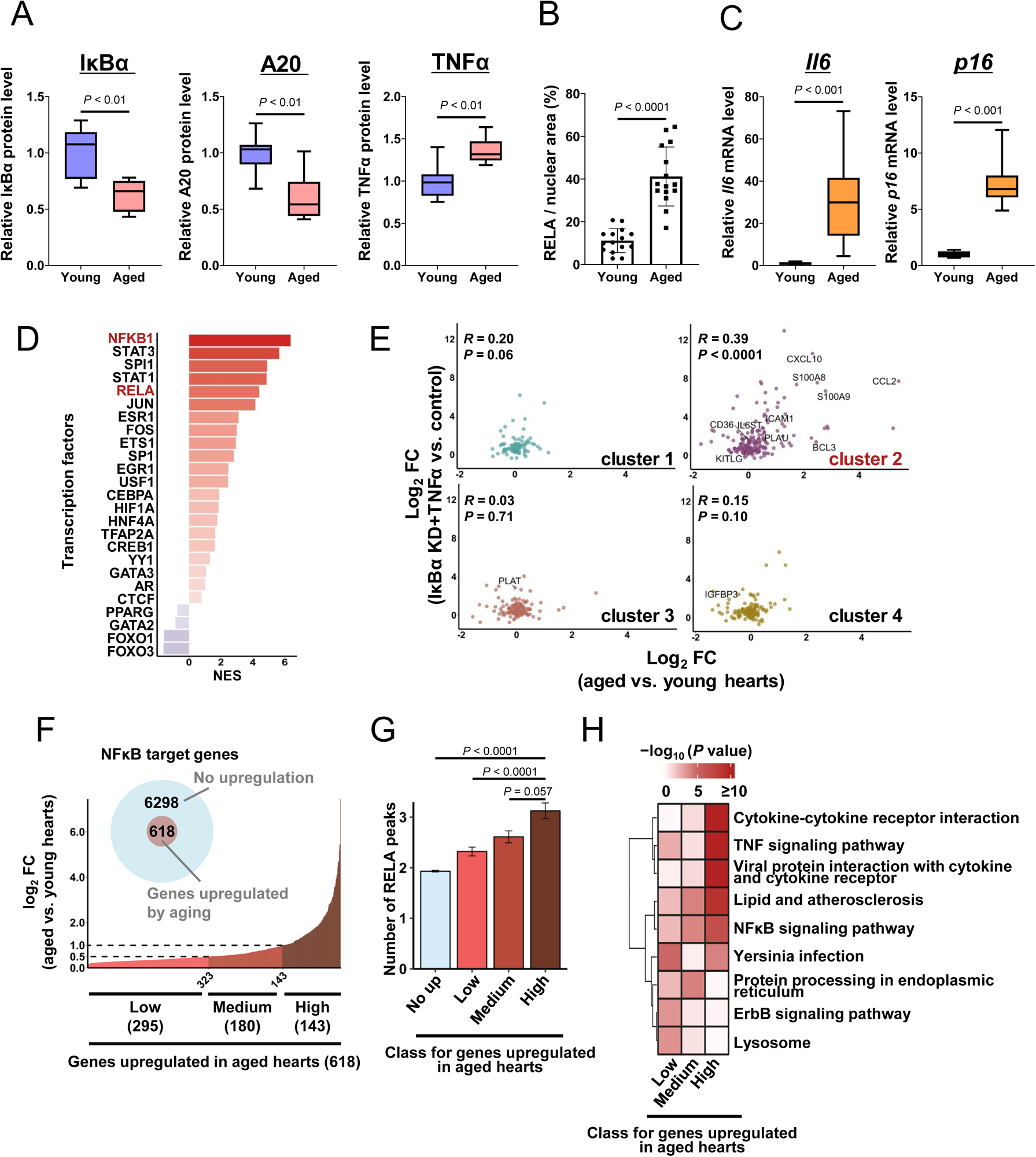
Sustained NFκB activation in the heart tissues of aged mice. (A) Protein expression of IκBα, A20, and TNFα in the hearts of young (8 weeks old, N = 8) and aged mice (69–73 weeks old, N = 8). (B) Immunohistochemistry of RELA in heart tissues from young and aged mice. Young mice: 8 weeks old, N = 3. Aged mice: 69–73 weeks old, N = 3. Bar plots showing the percentage of RELA positive staining relative to nuclear areas. Five HPFs were obtained from each heart tissue. (C) mRNA expression of *Il6* and *p16* in the hearts of young (8 weeks old, N = 5) and aged mice (69–73 weeks old, N = 5). (D) Transcription factor activity predicted by DorothEA. Confidence was set to A, and minsize was set to 30. (E) Correlation between the expression FC (IκBα KD+TNFα vs. Control MCF-7) of cluster 2 genes and FC of aged vs. young mouse hearts. Cellular senescence-promoting and SASP genes are labeled. (F) Classification of the NFκB target genes upregulated by aging in the mouse hearts. Light blue circle indicates NFκB target genes and pink circle indicates genes whose expression is markedly upregulated in aged mice (padj < 0.01, FC > 0). Genes included in the pink circle were classified according to the degree of gene expression with aging. (Low: 0 < FC ≤ 0.5, Medium: 0.5 < FC ≤ 1.0, High: 1.0 < FC). (G) Average count of RELA ChIP peaks annotated with genes in each group classified based on FC expression in aged mouse hearts. (H) KEGG enrichment analysis of the high group genes in aged hearts analysis in Figure 4F. Data represent the mean ± SD (B), ± SE (G), or median and interquartile range (25th and 75th percentiles), with error bars showing minimum to maximum [(A) and (C)]. *P* values were determined using the Kruskal– Wallis test followed by Steel–Dwass (G) or Mann–Whitney U test [(A), (B), and (C)]. The correlation was analyzed using the Spearman’s correlation (E).

### Sustained NFκB activity downregulates mTORC2 signaling

Next, we compared the changes in the signaling and downstream cascade between TNFα and TNFα+IκBα KD and the relationship with cellular senescence. First, we focused on the decreased expression of Cyclin D1 by IκBα KD+TNFα in MCF7 cells because it is related to the suppression of cell proliferation (Figure 1H). Cyclin D1 plays an important role in cell cycle entry at G1/S, and its expression and stability are regulated by multiple signaling pathways, including ErbB, MEK-ERK, PI3K-AKT, and NFκB ^34^. Therefore, we examined the effect of those pathway inhibitors (Figures 5A and S5) and found that AKT inhibitor VIII (AKTi-VIII; AKT1/2 inhibitor), SB203580 (inhibitor for p38 and AKT), and BAY-117085 (NFκB inhibitor) abrogated the changes in Cyclin D1 expression under IκBα KD+TNFα conditions (Figure 5A). The AKT and NFκB pathways are known to crosstalk with each other ^35, 36^. AKT is important for the translation and stability of the Cyclin D1 protein ^37^ and is also shown to be activated in G2 phase *via* the mTOR complex 2 (mTORC2) cascade ^38^; therefore, we analyzed the signaling pathway involved in AKT. As a result, we found that the expression of RICTOR, P-RICTOR, and P-AKT (Ser473) in the mTORC2 cascade was significantly reduced by IκBα KD+TNFα (Figure 5B). Moreover, AKTi-VIII treatment increased SA-β-gal positive cells and abolished IκBα KD+TNFα effects on SA-β-gal staining (Figure 5C). We also found that RICTOR KD increased SA-β-gal staining and decreased Cyclin D1 expression, as shown under the IκBα KD+TNFα condition (Figure 5D). These results suggested that sustained NFκB activity, which is mimicked by TNFα+IκBα KD, suppresses mTORC2/AKT signaling and Cyclin D1 expression, thereby contributing to promoting cellular senescence. Moreover, the MEK inhibitor U0126 partially abolished the effect of IκBα KD+TNFα on Cyclin D1 downregulation (Figure 5A), and IκBα KD+TNFα slightly reduced the P-ERK level (Figure 5B). Since ERK activity is important for inducing *CCND1* ^39^, downregulating the Cyclin D1 protein by IκBα KD+TNFα might be elicited *via* suppression of both the mTORC2/AKT and ERK signals and their crosstalk.

**Figure 5.**
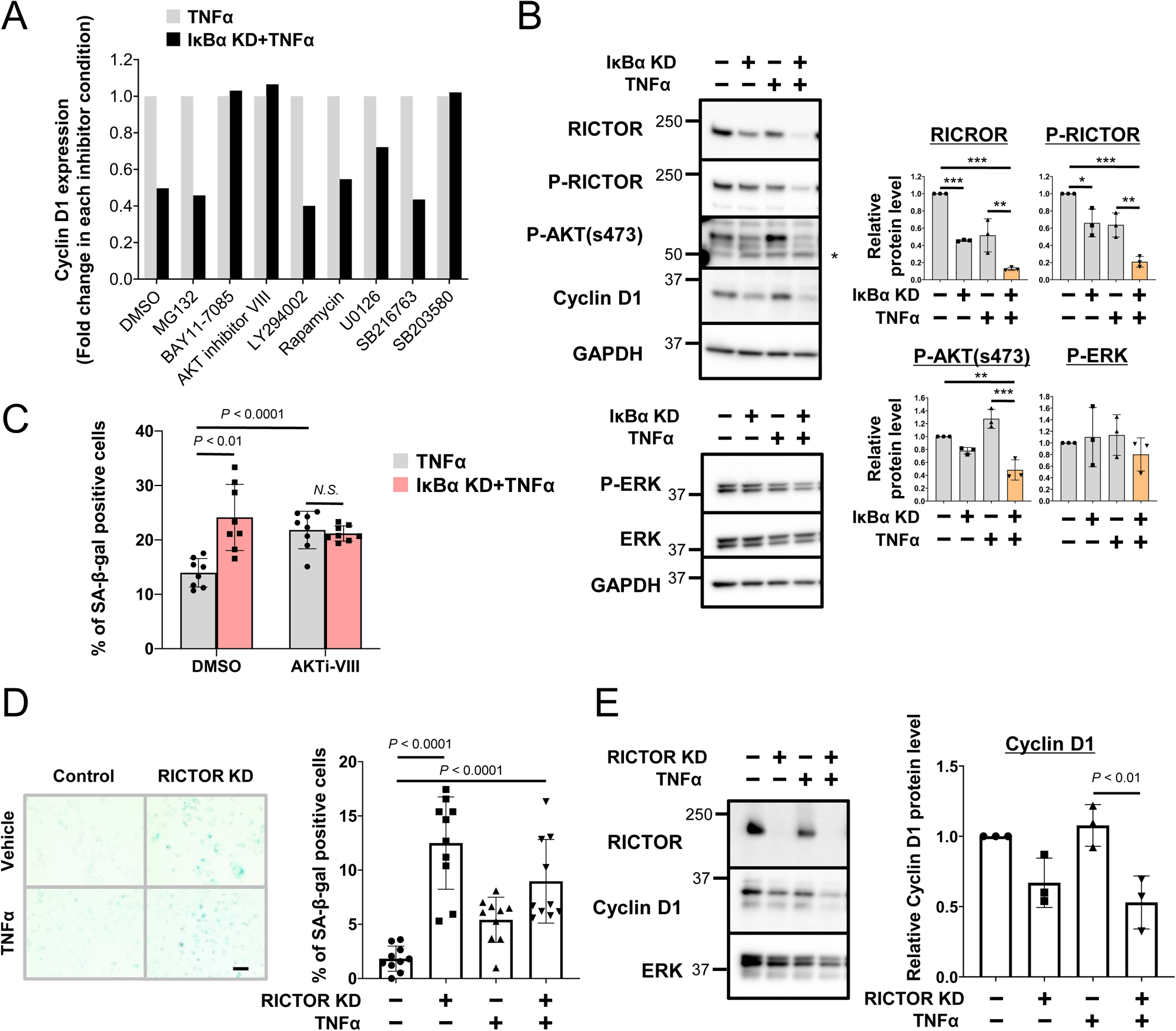
Effect of sustained NFκB activation in mTORC2 signaling. (A) Effect of AKT inhibitor on Cyclin D1 expression in MCF-7 cells under IκBαKD+TNFα conditions. Bar plots quantifying the relative Cyclin D1 level in each inhibitor condition. Cells were treated with BAY11-7085 (5 µM, NFκB inhibitor), AKTi VIII (1 µM, AKT1/2 inhibitor), LY294002 (10 µM, PI3K inhibitor), Rapamycin (100 nM, mTOR inhibitor), U01268 (10 µM, MEK inhibitor), SB216763 (10 µM, GSK3β inhibitor), SB203580 (10 µM, inhibitor for p38 and AKT), and TNFα for 48 hr. MG132 (10 μM, proteosome inhibitor) was added 8 hr before collecting samples. (B) Effect of IκBα KD+TNFα on protein expression of RICTOR, P-RICTOR, P-AKT(s473), and P-ERK in MCF-7 cells. Cells were treated with TNFα for 48 hr. (C) Effect of AKT inhibitor VIII (10 µM) on SA-β-gal staining in IκBα KD MCF-7 cells treated with TNFα. SA-β-gal staining in IκBα KD MCF-7 cells treated with TNFα and AKT inhibitor VIII (10 µM) for 24 hr. (D) Effect of RICTOR KD on SA-β-gal staining in MCF-7 cells. Right, representative images. Scale bars, 100 μm. Left, bar plots showing the percentage of SA-β-gal positive cells. (E) Effect of RICTOR KD on Cyclin D1 expression. Left, representative blot image. Right, bar plots showing quantification of relative protein levels of Cyclin D1. Data represent the mean ± SD [(B), (C), (D), and (E)]. *P* values were determined using one-way ANOVA followed by Tukey’s test [(B), (C), (D), and (E)]. \**P* < 0.05, \*\**P* < 0.01, \*\*\**P* < 0.001.

### Purine catabolism is induced by sustained NFκB activity

mTORC2 is a critical factor in metabolic regulation ^40^. Since metabolic dysregulation is a hallmark of cellular senescence and aging ^23^, we investigated metabolic alterations induced by sustained NFκB activation. We performed a metabolome analysis of MCF-7 cells under IκBα KD+TNFα conditions; however, principal component analysis showed similar intracellular metabolic profiles between TNFα and IκBα KD+TNFα (Figure S6A and Supplementary Data S1). TNFα alters several metabolic pathways, including glycolysis, lipid metabolism, and amino acid metabolism ^41–43^, which might mask metabolic changes associated with cellular senescence under IκBα KD+TNFα conditions. Since IκBα KD alone moderately reduced mTORC2 activity (Figure 5B), we focused on metabolic alterations induced by IκBα KD. The results showed that 62 and 35 metabolite levels were up- and downregulated, respectively (Figure 6A). Notably, the intracellular level of hypoxanthine, a potential oxygen generator ^44^, in the purine catabolic pathway was significantly elevated by IκBα KD (*P* < 0. 0001, FC = 3.126, vs. control), and its effects were enhanced by TNFα (*P* < 0. 0001, FC = 4.711, vs. control) (Figure 6B). The extracellular hypoxanthine level in the culture medium was also increased by IκBα KD+TNFα (*P* < 0. 0001, FC = 17.69, vs. control) (Figures S6B and S6C). Pathway enrichment analysis of metabolites altered by IκBα KD also confirmed the global changes in the purine metabolic pathway (Figure 6C). Mapping of intracellular metabolite changes induced by IκBα KD showed an increase in metabolites of the catabolic pathway of nucleotides (e.g., hypoxanthine, xanthine) and a decrease in high-energy molecules (e.g., ATP and GTP) (Figure 6D). Biosynthesis of purine nucleotides has been reported to increase the cellular growth rate and promote the G1/S transition ^45^. Moreover, an imbalance of nucleotide species suppresses cell proliferation, but it still causes a transition to S phase *via* DNA replication stress signaling ^46^. Therefore, our results suggest that IκBα KD cells inhibit normal cell cycle progression by inducing a nucleotide imbalance, including high-energy nucleotides.

**Figure 6.**
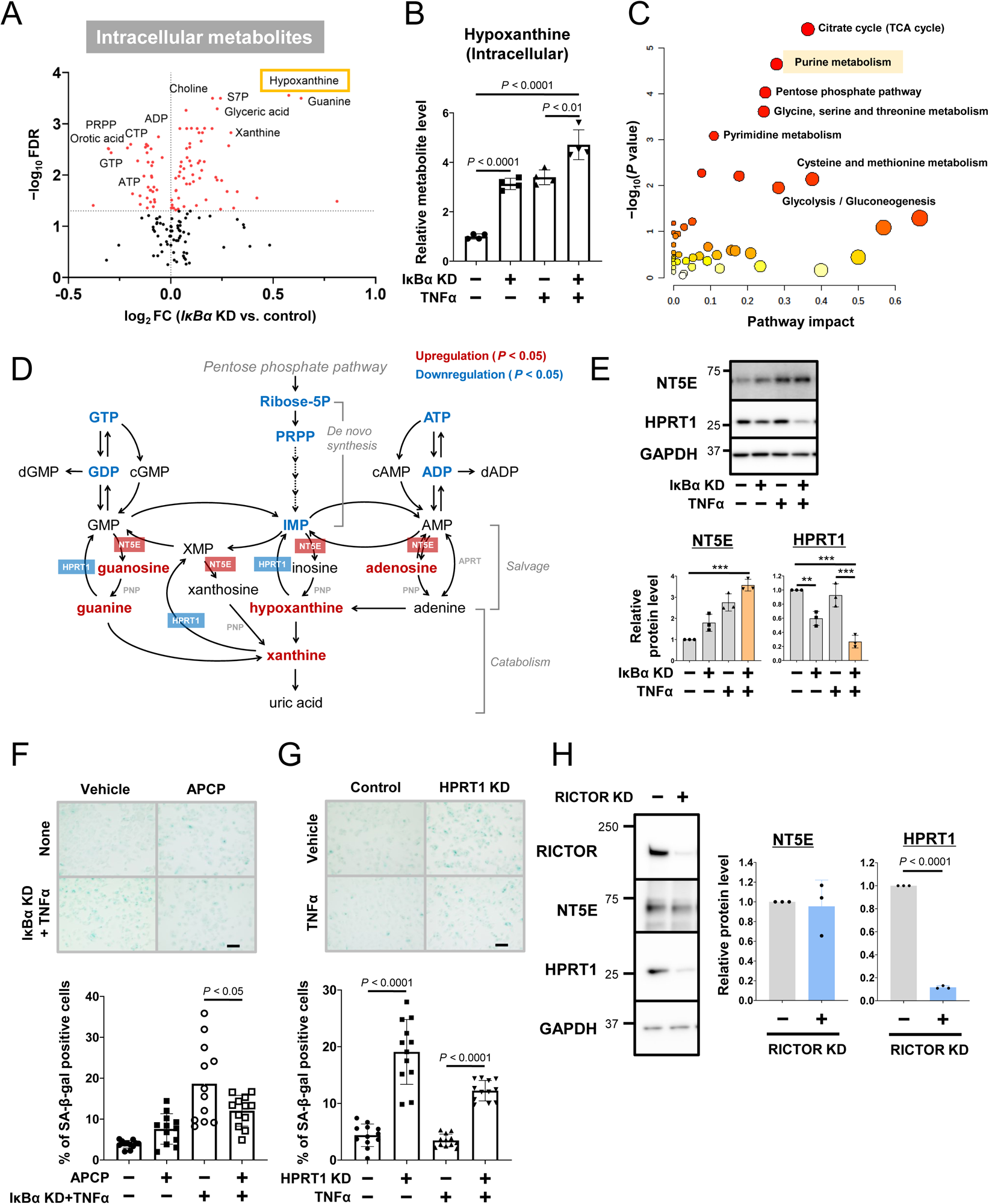
Purine catabolism induced by sustained NFκB activity. (A) Volcano plots showing differences in intracellular metabolites in MCF-7 cells under control and IκBα KD conditions. (B) Intracellular hypoxanthine levels in IκBα KD+TNFα. Cells were treated with TNFα for 48 hr. (C) Metabolite pathway enrichment analysis (MPEA) of intracellular metabolite differences between control and IκBα KD+TNFα. (D) Schematic purine metabolic pathway. Red indicates upregulation (*P* < 0.05), and blue indicates downregulation (*P* < 0.05). (E) Effect of IκBα KD on the protein expression of NT5E and HPRT1 in MCF-7 cells treated with TNFα for 48 hr. (F) Effect of NT5E inhibitor APCP (100 µM) on SA-β-gal staining in IκBα KD MCF-7 cells treated with TNFα. Cells were treated with APCP and TNFα for 24 hr. Upper, representative images. Scale bars, 100 μm. Lower, bar plot indicating the percentage of SA-β-gal positive cells. (G) Effect of HPRT1 KD on SA-β-gal staining in MCF-7 cells treated with TNFα for 24 hr. (H) Effect of RICTOR knockdown on protein expression of NT5E and HPRT1 in MCF-7 cells. Cells were treated with TNFα for 48 hr. Data represent the mean ± SD [(B), (E), (F), (G), and (H)]. *P* values were determined using unpaired two-tailed Student’s t tests (H) or one-way ANOVA followed by Tukey’s test [(B), (E), (F), and (G)]. \*\**P* < 0.01, \*\*\**P* < 0.001.

More precisely, a balance between the synthesis and degradation of purine nucleotides determines the cellular level of adenylate and guanylate ^45^. Two pathways have been identified for purine nucleotides synthesis: the *de novo* pathway and salvage pathway ^47^. Detailed analysis of mRNA and protein levels in IκBα KD cells validated the promotion of purine catabolism, including the increased ecto-5’-nucleotidase (NT5E) level and decreased hypoxanthine phosphoribosyltransferase (HPRT) 1 level, which were enhanced by TNFα (Figures 6E and S6D). In the purine salvage pathway, NT5E catalyzes the hydrolysis of adenosine monophosphate (AMP) to adenosine ^48^, and HPRT1 catalyzes the conversion of hypoxanthine and guanine to inosine monophosphate (IMP) and monophosphate (GMP), respectively ^49^ Notably, several other genes and enzymes related to purine *de novo* synthesis were also downregulated by IκBα KD+TNFα (Figure S6D). To address whether upregulation of purine catabolism is one of the causes of the progression of cellular senescence, we next tested the effect of NT5E and HPRT1 inhibition. As predicted, the NT5E inhibitor adenosine 5’-(α, β-methylene) diphosphate (APCP) attenuated the increase in SA-β-gal positive cells by IκBα KD+TNFα while siRNA-mediated HPRT1 KD augmented SA-β-gal staining (Figures 6F, 6G, and S6E). Furthermore, RICTOR knockdown decreased HPRT1 expression but did not affect NT5E expression (Figure 6H), indicating that HPRT1 expression is controlled by RICTOR. MCF-7 cells under the IκBα KD+TNFα condition downregulated Cyclin D1 expression and prolonged the cell cycle (Figures 1H and 1I). This may be due to the depletion of nucleotides and high-energy metabolites, such as ATP, GTP, dATP, and dGTP, which are necessary for protein translation and cell cycle progression ^47^.

Next, we examined whether similar changes in purine catabolism mediated by NFκB pathway are also observed in the hearts of aged mice *in vivo*. Protein expression of RICTOR and HPRT in aged hearts was significantly lower than that in young hearts (Figures 7A 7B, and S7A). The protein levels of RICTOR (R = 0.7676, *P* < 0.001) and HPRT (R = 0.7676, *P* < 0.001) were highly correlated with the IκBα levels in the hearts of individual young and aged mice, respectively (Figures 7A, right and 7B, right). Metabolome analysis of the hearts of aged mice also showed enhanced purine catabolic metabolism (xanthosine, xanthine, and GMP) and reduced high-energy molecules (ATP and ADP) (Figures 7C, S7B, and Supplementary Data 2). Interestingly, mass spectrometry imaging revealed that ATP levels in the hearts of aged mice were markedly reduced in the entire tissue (Figures 7D and S7C). As the aged heart tissues exhibited a higher amount of NFκB and lower amount of IκBα (Figures 4A and 4B), these metabolic features presumably represent a mechanism underlying the ability of highly nuclear-localized NFκB to interfere with the synthesis of high-energy metabolites.

**Figure 7.**
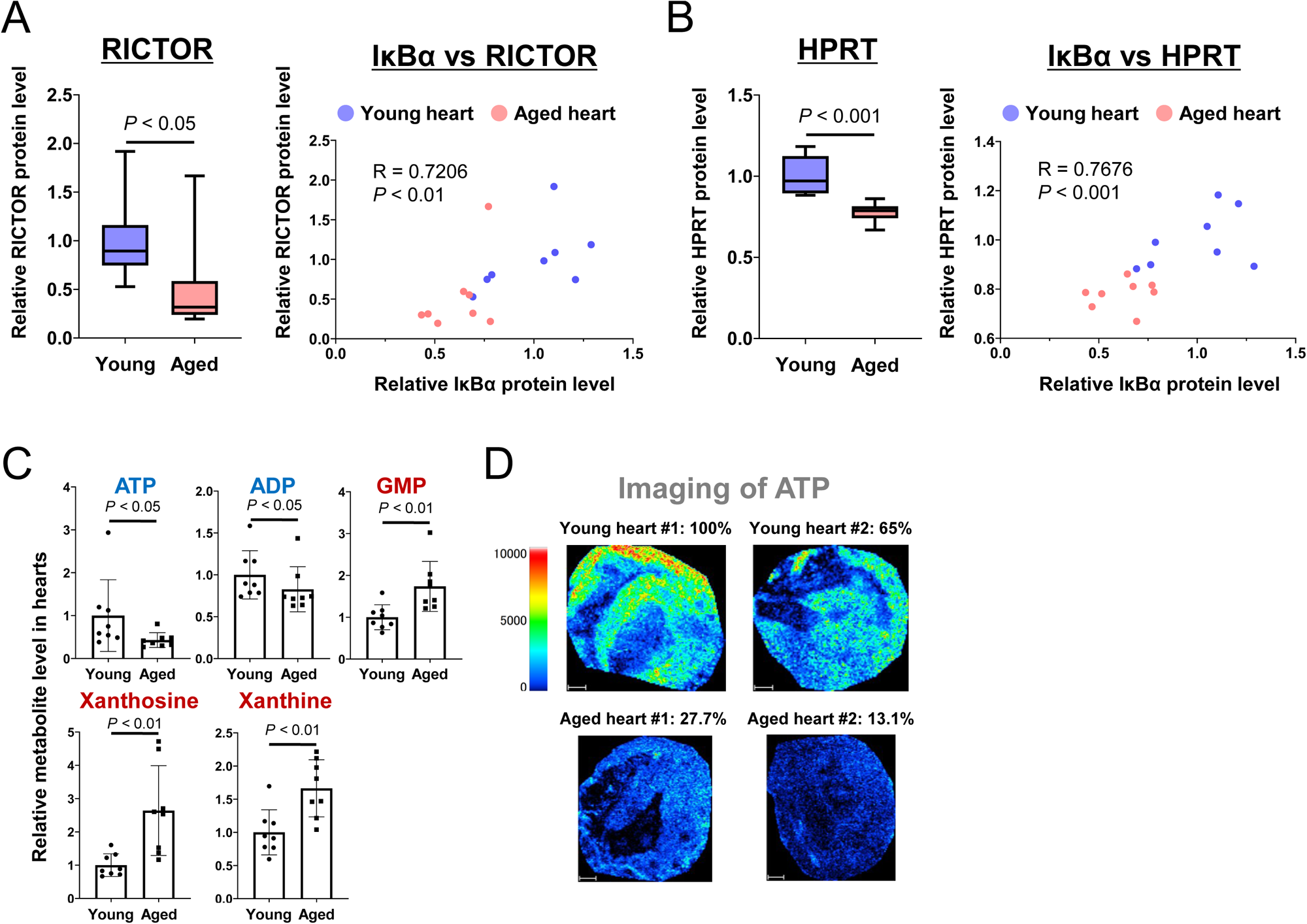
Purine catabolism in the heart tissues of aged mice. (A and B) Protein expression of RICTOR (A) and HPRT1 (B) in young and aged hearts. Correlations between RICTOR and IκBα protein expression in young and aging heart tissue (A, right) and HPRT and IκBα protein expression (B, right). (C) Metabolites significantly changed in the purine metabolic pathway between young and aged heart tissues. Young mice: 8 weeks old, N = 8. Aged mice: 69–73 weeks old, N = 8. (D) Mass spectrometric images of ATP levels in young and aged heart tissues. The percentage shows the ATP level relative to young heart #1. Upper, young hearts; lower, aged hearts. Data represent the mean ± SD (C) or median and interquartile range (25th and 75th percentiles) with error bars showing minimum to maximum [(A) and (B)]. *P* values were determined using Mann–Whitney U test [(A), (B), and (C)]. The correlation was analyzed using the Spearman’s correlation [(A) and (B)].

## DISCUSSION

The duration of nuclear NFκB activity has various effects on gene expression by regulating mRNA half-life or epigenetic status ^20, 21^. In this study, we show that the aging-associated phenotypic changes could be explained by a core regulatory mechanism stemming from sustained nuclear NFκB activity. That is, increased nuclear NFκB enhances NFκB-DNA binding for inflammatory gene expression, alters the mTORC2/AKT pathway to promote purine catabolism, and inhibits cell cycle progression.

Our study showed that the loss of two negative feedback regulators, namely, IκBα and A20, significantly elevated the nuclear abundance of NFκB and induction of cell senescence. However, a combination of IκBα KD and A20 OE did not strongly alter the NFκB dynamics from sustained to oscillatory but rather reduced the amplitude of nuclear NFκB activity and the expression of *IL8* and *CCL2* genes, which were upregulated by IκBα KD alone (Figures 2C and 2H). These results suggest that the nuclear NFκB level is critical for inflammatory aging. Indeed, the decrease in IκBα protein levels under oxidative stress in MCF-7 cells (Figure S7D) indicates that decreases in IκBα may occur over a lifetime for the same reason. Meanwhile, previous studies have demonstrated that defects in the feedback regulation of IκBα in the NFκB pathway cause Sjögren’s syndrome and shortened life span ^50^ and that mice expressing vascular-specific IκBα overexpression have a prolonged life span ^51^. However, further studies are needed to determine the mechanism between sustained NFκB activity and its effect on life span.

Notably, inhibition of purine nucleoside phosphorylase, which is one of the rate-limiting enzymes for purine catabolism, suppresses cellular senescence *in vivo* ^52^ and promotes the synthesis of nicotinamide adenine dinucleotide, an anti-aging molecule ^53^. On the other hand, hypoxanthine, a purine catabolic metabolite and a common DNA damage base that has mutagenic potential due to its function as a potential radical generator ^44, 54^, was significantly elevated under the IκBα KD+TNFα condition. Therefore, hypoxanthine may be a key molecule in NFκB-associated cell senescence and likely contributes to accelerating the aging process.

Our study showed that the cell cycle period is prolonged during the G2/M phase under IκBα KD+TNFα. However, issues regarding the G2/M delay mediated by metabolic imbalance remain unresolved. Although the shortage of high-energy metabolites and nucleotides severely affect the G1/S transition, the effect of downregulating purine metabolism at the G2/M phase was not clear ^45^. An earlier study showed that NFκB activation in response to ionizing radiation or etoposide arrested cells at the G2/M phase for a prolonged time ^55^. Since hypoxanthine potentially generates free radicals, G2/M cell cycle arrest under IκBα KD+TNFα may involve hypoxanthine upregulation. In addition, mTORC2 and AKT activity are required for G2/M cell cycle progression ^38, 55^, and our data show that RICTOR, P-RICTOR, and P-AKT (Ser473) expression in the mTORC2 cascade was significantly reduced by IκBα KD+TNFα. An imbalance of nucleotide species has been reported to suppress cell proliferation, although the cells still pass through S phase by the activation of DNA replication stress signaling ^46^. Therefore, inhibiting or downregulating the cell cycle speed under IκBα KD+TNFα may be caused by either upregulation of hypoxanthine *via* HPRT1 and mTORC2 or downregulation of purine metabolism for high-energy metabolite synthesis. Whether another mechanism underlies the cell cycle regulation observed in this study requires further investigation. Identifying a mechanism that can distinguish between cellular senescence and cell death induced by TNFα-dependent CDK regulation would be also important in future study ^56^. Overall, in this study, we show that temporal NFκB dynamics affect an unexpectedly broad spectrum of cell physiological factors and regulate cellular senescence. These findings provide unprecedented insights into the molecular basis of cellular senescence.

## Supporting information

Key Resources Table

## ACKNOWLEDGMENTS

We thank Dr. Keita Iida, Dr. Hiroaki Imoto, Mr. Ken Murakami, and Mr. Masatoshi Haga for engaging in discussions regarding computational analysis and performing critical readings of the manuscript. We also thank Ms. Emiko Suga, Ms. Akimi Mizoguchi, Mr. Hiroki Michida, and Mr. Johannes Nicolaus Wibisana for their help with the mouse experiments, qRT-PCR, western blot analysis, and plasmid construction. We also thank Dr. Kazuhiro Aoki for the generous gift of the Fucci probe, Mr. Hideya Aragaki from Olympus Corporation for providing the cell tracking software, and Dr. Shigeyuki Magi and Dr. Yutaka Suzuki for their help with the RELA ChIP-seq analysis. We also thank Ms. Maiko Goto and Ms. Nanako Seki for their help with the metabolome analysis. This study was supported by the JSPS KAKENHI (17H06299, M.O.; 17H06302, M.O.; 18H04031, M.O.; JP17H06304; T.B.; JP17H06299, T.B.), the JST CREST Program (JPMJCR21N3, M.O.), the Uehara Memorial Foundation (M.O.), the AMED BINDS Program (JP23ama121055. T.B.), and the Cooperative Research Project Program of the Medical Institute of Bioregulation, Kyushu University (S.T. and T.B.).

## AUTHOR CONTRIBUTIONS

Conceptualization: S.T., K. Matsuda, and M.O.; Formal analysis: S.T., K. Matsuda, and K.N.; Funding acquisition: T.B. and M.O.; Investigation: S.T., Y.I., M.T., Y.M., A.I.N., and S.S.; Methodology: S.T., K. Matsuda, K.N., Y.I., M.T., Y.M., A.I.N., S.S., K. Moro, T.B., and M.O.; Project administration: S.T. and M.O.; Resources: K. Moro, T.B., and M.O.; Supervision: K. Moro, T.B., and M.O.; Visualization: S.T., K. Matsuda, K.N., A.I.N., and M.O.; Writing: S.T., K. Matsuda, K.N., A.I.N., and M.O.

## STAR METHODS

### EXPERIMENTAL MODEL AND SUBJECT DETAILS

#### Reagents and cell culture

TNFα was purchased from Thermo Fisher Scientific (Waltham, MA, USA). MG132, Rapamycin, and LY294002 were purchased from Calbiochem (Merck Millipore, Burlington, MA, USA), BAY11-7085 was purchased from Fujifilm Wako Pure Chemical Industries, Ltd. (Osaka, Japan). AKT inhibitor VIII was purchased from Cayman Chemical (Ann Arbor, MI, USA). SB216763 was purchased from Sigma-Aldrich (St Louis, MO, USA). SB203580 was purchased from Cell Signaling Technology (Danvers, MA, USA). U0126 was purchased from MedChemExpress (Monmouth Junction, NJ, USA). APCP was purchased from Tocris Bioscience (Bristol, UK). Methanol, Hydrochloric Acid, Ethanol, Xylene, Mayer’s hematoxylin solution, 1% Eosin Y solution, and PathoMount were purchased from Fujifilm Wako Pure Chemical Industries, Ltd. 9-Aminoacridine (9AA) was purchased from Merck (Darmstadt, Germany). Polyethylene Glycol 600 Sulfate for MALDI-MSI was purchased from Tokyo chemical industry (Tokyo, Japan). Indium tin oxide (ITO)-coated glass slides (100 Ω/sq without adhesive material coating) for MALDI-MSI and adhesive-coated glass slides for hematoxylin and eosin staining (HE), and coverslips were purchased from Matsunami Glass (Osaka, Japan).

The human breast adenocarcinoma MCF-7 cell line was obtained from the American Type Culture Collection (ATCC; Manassas, VA, USA). Cells were grown in Dulbecco’s modified Eagle’s medium (Nakalai Tesque, Kyoto, Japan) containing 10% (v/v) fetal bovine serum (FBS; Sigma-Aldrich), and antibiotics (100 U/mL penicillin, 100 mg/mL streptomycin, and 0.25 mg/mL amphotericin B; Nakalai Tesque, Kyoto, Japan) at 37 °C under a humidified atmosphere with 5% CO_2_.

#### Mouse experiments

All mice were purchased from the Japan SLC and maintained under specific pathogen–free conditions at Graduate School of Frontier Biosciences Osaka University. Female 8-weeks-old (young) and 69–73-weeks-old (aged) C57BL/6 N mice were used in experiments. All animals were separately caged with a 12:12-h light–dark cycle and had free access to water and chow throughout the study. The excised tissues were immediately frozen in liquid nitrogen and stored at −80°C until use. All experiments were approved by the Animal Care and Use Committee of Osaka University and were performed following institutional guidelines.

## METHODS

### Quantitative RT-PCR (qRT-PCR) analysis

Real-time PCR analysis was conducted as described previously ^57^. Briefly, RNA was isolated from cells or tissues using NucleoSpin RNA Plus (Macherey-Nagel, Duren, Germany, Cat. No. 740984.250) and subjected to cDNA synthesis using ReverTra Ace qPCR RT Master Mix (TOYOBO, Osaka, Japan) and quantitative PCR using a KOD SYBR qPCR kit (TOYOBO) with CFX96 Real-Time PCR System (Bio-Rad Laboratories, Hercules, CA, USA) according to the manufacturer’s protocol. The ΔΔCq method was used to quantify gene expression, using RPL27 or Rplp0 expression as an internal reference ^58^. All experiments were performed in triplicate. The primers used for real-time PCR are listed in Supplementary Table S4.

### Western blotting

Immunoblot analysis was performed as described previously ^57^ using primary antibodies against GAPDH (1:2000; Sigma-Aldrich, MAB374), GAPDH (1:4000; Proteintech, MA, USA, 10494-1-AP), αTubulin (1:1000; Abcam, Cambridge, MA, USA, ab15246), IκBα (1:1000; Cell Signaling Technology, Danvers, MA, USA, #4814), IκBα (1:1000; Cell Signaling Technology, #4812), RELA (1:1000; Cell Signaling Technology, #8242), A20 (1:1000; Cell Signaling Technology, #5630), RICTOR (1:500; Santa Cruz Biotechnology, Inc., CA, USA, sc-81538), RICTOR (1:1000; Cell Signaling Technology, #5379), P-RICTOR (1:1000; Cell Signaling Technology, #3806), AKT (1:1000; Cell Signaling Technology, #4685), P-AKT(Ser473) (1:1000; Cell Signaling Technology, #4051), P-AKT(Ser473) (1:1000; Cell Signaling Technology, #4060), P-AKT(Thr308) (1:1000; Cell Signaling Technology, #2965), NT5E (1:1000; Cell Signaling Technology, #13160), HPRT1 (1:1000; Proteintech, 15059-1-AP), HPRT1 (1:500; Santa Cruz Biotechnology, Inc., sc-376938), Cyclin D1 (1:1000; Cell Signaling Technology, #2978), IKKβ (1:1000; Cell Signaling Technology, #2370), P-IKKα/β (Ser176/180) (1:1000; Cell Signaling Technology, #2697), Lamin A (1:1000; Abcam, ab8980), ERK (1:1000; Cell Signaling Technology, #9102), P-ERK (1:1000; Cell Signaling Technology, #4370), GFP (1:500; Santa Cruz Biotechnology, Inc., sc-5385), and secondary antibodies [HRP-linked anti-mouse IgG (1:5000; Cell Signaling Technology, #7076), HRP-linked anti-goat IgG (1:5000; Santa Cruz Biotechnology, Inc., sc-2020), and HRP-linked anti-rabbit IgG (1:5000; Cell Signaling Technology, #7074)].

For the separation of nuclear fraction, the nuclear and cytoplasmic protein extraction kit (Biovision Inc., Milpitas Boulevard, Milpitas, CA, USA, K266) was used according to the manufacturer’s procedures.

### RELA immunocytochemistry

Cells were fixed with 4% paraformaldehyde (Thermo Fisher Scientific) in phosphate buffered saline (PBS) for 15 min, rinsed with PBS, and permeabilized with 0.2% Triton X-100 (Nakalai Tesque) in PBS for 10 min. Cells were blocked using 1% goat serum Thermo Fisher Scientific, #16210–064) in PBS for 15 min and incubated with RELA antibodies (1:1600; Cell Signaling Technology, #8242) diluted in 1% goat serum/PBS at 4 °C overnight. After rinsing in PBS, cells were incubated with Alexa Fluor 488–conjugated secondary antibodies (1:2000; Thermo Fisher Scientific, A32731), 0.125 ng/mL CellMask^TM^ (Thermo Fisher Scientific) as a cell body stain, and 1 µg/mL 4’,6-diamidino-2-phenylindole (DAPI; Nacalai Tesque) as a nuclear stain for 1 h at 25 °C in the dark. Fluorescent images were acquired using IN Cell Analyzer 2500HS (Cytiva, Preston, UK) at 20× magnification. Images of 30 fields were taken for each condition. CellProfiler (ver. 3. 1. 9) was used to segment cellular regions from CellMask^TM^ images, to segment nuclear regions from DAPI images, and to quantify RELA signal intensities for each cell ^59^. The nuclear-to-cytoplasm signal ratio for RELA was calculated based on the integrated signal density.

### Analysis of cell morphology

Cell morphology analysis was conducted as described previously ^57^. Briefly, Cells were fixed using 4 % paraformaldehyde for 15 min and permeabilized with 0.1 % Triton-X100 for 5 min. Cell membrane and nucleus were stained with 0.125 ng/mL CellMask^TM^ (Thermo Fisher Scientific) and 1 μg/mL DAPI (Nacalai Tesque) for 30 min at 25 °C in the dark, respectively. Fluorescent images were acquired using IN Cell Analyzer 2500HS (Cytiva, Preston, UK) at 20× magnification. CellProfiler (ver. 3. 1. 9) was used to extract geometrical features from the cell images ^59^. DAPI and CellMask^TM^ staining was used to visualize the segmentation of nucleus and cell body, respectively. The following morphological parameters of the cells were evaluated. Area, Form Factor, Solidity, Extent, Orientation, Eccentricity, Compactness, major axis length, minor axis length, maximum ferret diameter, minimum ferret diameter, Perimeter, and Mean radius (total 13 parameters). PCA based on the 13 parameters was performed by the “prcomp” function in the “stats” R package.

### Senescence associated β-galactosidase (SA-β-gal) staining

The senescent cells were stained by a senescence β-galactosidase staining Kit (#9860, Cell Signaling Technology), according to the manufacturer’s instructions. Briefly, cells in 12-well plates were washed with PBS, fixed for 10 min in a fixative solution, washed twice with PBS, and incubated in SA-β-gal staining solution overnight at 37 °C. Images were acquired with a Keyence BZ-9000 microscope on 20× magnification. Four random views per well were selected to count the number of cells stained positive for SA-β-gal. Each independent experiment was repeated three times and the percentage of SA-β-gal positive cells per field from a total of 12 fields was analyzed with Image J software. DAPI staining was used to identify nuclei and count the number of cells per field. The threshold for identifying senescent cells was set by a positive control (etoposide treatment).

### siRNA transfection

siRNA duplexes for IκBα (ON-TARGET plus SMART pool, L-004765-00), RELA (ON-TARGET plus SMART pool, L-003533-00), A20 (ON-TARGET plus SMART pool, L-009919-00), RICTOR (ON-TARGET plus SMART pool, L-016984-00), HPRT1 (ON-TARGET plus SMART pool, L-008735-00), and negative control (D-001810-02) were purchased from Dharmacon/Horizon Discovery (Cambridge, UK). Cells were transfected with siRNA oligomer (final concentration 50 nM) mixed with Lipofectamine RNAiMAX reagent (Thermo Fisher Scientific) in serum-reduced Opti-MEM (Thermo Fisher Scientific) according to the manufacturer’s instructions.

### Cell-viability assay

Cell viability was measured using trypan blue exclusion assay. A 2 × 10^5^ cells/well was seeded in 12-well plates and incubated at 37 °C. After treatment with reagents and/or siRNAs, cells were disaggregated in 400 μL medium, and 10 μL of the suspension was mixed with 10 μL trypan blue (Thermo Fisher Scientific). Viable cells were counted using a Countess Automated Cell Counter (Thermo Fisher Scientific).

### Cell cycle analysis using the FUCCI biosensor

MCF-7 cells stably expressing the FUCCI biosensor were obtained by the PiggyBac transposase system ^60^. MCF-7 cells were transfected with the pPBbsr2-H2B-iRFP-P2A-mScarlet-I-hGem-P2A-PIP-tag-NLS-mNeonGreen (https://benchling.com/s/seq-LPZ1tLdpgnpJIYOG2ujR) and the hyPBase (https://benchling.com/s/seq-oGkw53b41IZqvzF5yQ9K) ^61^ plasmids using Lipofectamine LTX following manufacturer’s protocol (Thermo Fisher Scientific). The plasmids, pPBbsr2-H2B-iRFP-P2A-mScarlet-I-hGem-P2A-PIP-tag-NLS-mNeonGreen and hyPBase, were a gift from Dr. K. Aoki (National Institute for Basic Biology, National Institutes of Natural Sciences). After transfection, cells were released into a drug-free medium for 48 hr followed by blasticidin (10 µg/mL) selection until single colonies were formed. Single clones were expanded, and the FUCCI biosensor signal was confirmed by fluorescent microscopy. Time-lapse microscopy images were acquired using an IN Cell Analyzer 2500HS (GE Healthcare Life Science) with a CFI S Plan Fluor ELWD dry objective lens 20× (NA: 0.45) using an excitation wavelength of 475 nm and emission wavelength of 511 nm for PIP-tag-mNeonGreen and excitation wavelength of 542 nm and emission wavelength of 597 nm for mScarlet-hGem. Cells were maintained at 37 °C in a humidified atmosphere of 5% CO_2_ and imaged at 20 min intervals for 60 hr. Image processing and quantifications for a percentage of cells in each cell cycle phase were performed using tools in ImageJ-Fiji software (version 1.53t, NIH). Masks of mScarlet-hGem and PIP-tag-mNeonGreen signals were obtained by applying a fixed threshold (160, 65535). The obtained masks were identified with ‘Analyze Particles’ (size = 10.00–infinity, circularity = 0.00–1.00) and automatically counted as ROIs (region of interest). Cell cycle phases were determined by G1-phase: PIP-tag positive, S-phase: hGem positive, G2/M-phase: PIP-tag and hGem positive. The average fluorescence brightness of mScarlet and mNeonGreen was obtained by cell tracking software (Olympus). The threshold parameters of mScarlet and mNeonGreen for each cell were manually set. Subsequently, an algorithm for estimating the cell cycle phase assembled with R packages [brunnermunzel (v.1.4.1), multimode (v.1.4), rlist (v.0.4.6.1), tidyverse (v.1.3.0)] was applied to determine each cell cycle phase at the single-cell level. For quantifications of the G2/M phase length from single-cell data, measurements within 24–60 hr were used when the effects of TNFα were observed.

### Generation of IκBα KO MCF-7 cells

IκBα knockout (KO) MCF-7 cells were generated by CRISPR-Cas9 technology. We designed three independent guide RNAs targeting IκBα (NFKBIA) with the gRNA design tool (https://chopchop.cbu.uib.no/) ^62^. Two gRNAs were designed to target exon 1 and one gRNA to target intron 1 of NFKBIA (Supplementary Table S5). Primers were then designed by adding guanosine base on the 5’ and sticky ends corresponding to BbsI site. The pair of primers were then annealed and ligated into BbsI-digested eCas9-P2A-Puro plasmid ^63^. MCF-7 cells were transfected with IκBα gRNA-containing eCas9-P2A-Puro plasmids using Lipofectamine LTX following the protocol of the manufacturer (Thermo Fisher Scientific). After transfection, cells were released into a drug-free medium for 48 h followed by puromycin (1 µg/mL) selection until single colonies were formed. Single clones were expanded, and gene deletion was confirmed by western blotting. eSpCas9(1.1)-T2A-Puro was a gift from Andrea Németh (Addgene, # 101039).

### Generation of A20-overexpressing MCF-7 cells

To establish cell lines stably overexpressing A20, the PiggyBac transposase system was used ^60^. The plasmid used for A20 overexpression, pPB[Exp]-EGFP/Puro-CMV>hTNFAIP3, was constructed with VectorBuilder (Kanagawa, Japan). The vector ID is VB220607-1391zue and more information can be obtained from vectorbuilder.com. MCF-7 cells were transfected with the pPB[Exp]-EGFP/Puro-CMV>hTNFAIP3 and hyPBase ^61^ plasmids using Lipofectamine LTX following the manufacturer’s protocol. (Thermo Fisher Scientific). After transfection, cells were released into drug-free medium for 48 h followed by puromycin (1 µg/mL) selection until single colonies were formed. Single clones were expanded, and expression of A20 was confirmed by western blotting.

### Immunohistochemistry of RELA in heart tissue from aged mice

Immunohistochemistry was performed at Genostaff (Tokyo, Japan). Tissue sections were fixed with 4% paraformaldehyde for 10 min. Endogenous peroxidase was blocked with 0.3% H_2_O_2_ in methanol for 30 min, followed by incubation with G-Block (Genostaff) and avidin/biotin blocking kit (Vector Laboratories, Peterborough, UK). The sections were incubated with anti-REL A rabbit monoclonal antibody (0.004 µg/mL, Cell Signaling Technology, #8242) at 4 L overnight and then incubated with biotin-conjugated anti-rabbit IgG (Vector Laboratories) for 30 min at room temperature. Subsequently, peroxidase-conjugated streptavidin (Nichirei, Tokyo, Japan) was added for 5 min, and peroxidase activity was visualized using diaminobenzidine. The sections were counterstained with Mayer’s Hematoxylin (Muto Pure Chemicals Co., Ltd., Tokyo, Japan), dehydrated, and then mounted with Malinol (Muto Pure Chemicals Co., Ltd.).

Stained slides were digitally scanned using a NanozoomerS210 (Hamamatsu Photonics, Hamamatsu, Japan). For histopathological assessment, five random digital images of each mouse heart tissue were captured at 40× magnification using NDP view2 software (Hamamatsu Photonics). RELA-positive staining in nuclei was quantified by the ImmunotRatio ImageJ plugin ^64^. The brown (RELA) and blue (hematoxylin) threshold adjustments were set at 10 and 15, respectively.

### Metabolome analysis

Metabolome analysis was conducted as described previously ^65^. Briefly, the cell samples (approximately 1 × 10^6^ cells), conditioned media (50 µL), and heart tissues (approximately 30 mg) were mixed with 1 mL of cold methanol containing 10-camphorsulfonic acid (1.5 nmol) and piperazine-1,4-bis (2-ethanesulfonic acid) (1.5 nmol) as internal standards. Each sample was vigorously mixed by vortexing for 1 min, followed by 5 min of sonication. The extracts were then centrifuged at 16,000 × g for 5 min at 4 °C, and the resultant supernatant (400 μL) was collected. Protein concentrations in the pellet were determined using a Pierce™ BCA Protein Assay Kit (Thermo Fisher Scientific, MA). After mixing 400 μL of supernatant with 400 μL of chloroform and 320 μL of water, the aqueous and organic layers were separated by vortexing and subsequent centrifugation at 16,000 × g and 4 °C for 5 min. The aqueous (upper) layer (500 μL) was transferred into a clean tube. After the aqueous layer extracts were evaporated under vacuum, the dried extracts were stored at −80 °C until the analysis of hydrophilic metabolites. Before analysis, the dried aqueous layer was reconstituted in 50 μL of water.

Two liquid chromatography high-resolution tandem mass spectrometry (LC/MS/MS) methods for hydrophilic metabolite analysis were employed ^65, 66^. Anionic polar metabolites (i.e., organic acids, sugar phosphates, nucleotides, etc.) were analyzed via ion chromatography (Dionex ICS-5000+ HPIC system, Thermo Fisher Scientific) with a Dionex IonPac AS11-HC-4 μm column (2 μm i.d. × 250 mm, 4 μm particle size, Thermo Fisher Scientific) coupled with a Q Exactive high-performance benchtop quadrupole Orbitrap mass spectrometer (Thermo Fisher Scientific) (IC/MS/MS). Cationic polar metabolites (i.e., amino acids, bases, nucleosides, etc.) were analyzed via liquid chromatography (Nexera X2 UHPLC system, Shimadzu) with a Discovery HS F5 column (2.1 mm i.d. × 150 mm, 3 μm particle size, Merck) coupled with a Q Exactive instrument (PFPP-LC/MS/MS). The two analytical platforms for hydrophilic metabolite analysis were controlled using LabSolutions, version 5.80 (Shimadzu) and Xcalibur 4.2.47 (Thermo Fisher Scientific). Metabolome analyses and data processing were performed at the Division of Metabolomics of the Medical Institute of Bioregulation at Kyushu University. Quantitative levels of metabolites were calculated using peak areas relative to an internal standard (10-camphorsulfonic) and corrected for the total protein amount (analysis for MCF7 cells) or weight (analysis for mouse heart tissues) of each sample.

To interpret the data, the levels of the metabolites significantly altered by IκBα KD (FDR < 0.05) or aging in the mouse heart (FDR < 0.1) were analyzed for pathway enrichment using the web-based MetaboAnalyst 5.0 software (http://www.metaboanalyst.ca) and KEGG library (http://www.genome.jp/kegg/). Metabolite identifiers (Human Metabolome Database [HMDB] ID, http://www.hmdb.ca) were used for each metabolite.

### Analysis for ATP imaging in the mouse heart using MALDI-MSI Standard sample analysis

To detect ATP, 9AA was used as a matrix. The 9AA was dissolved in distilled water and methanol (3:7, v/v) at a concentration of 5 mg/mL. Each concentration standard solution and matrix solution were equivalent volume mixed. The mixture was spotted on ITO-coated glass slides, dried, and measured.

### MALDI-MSI analysis

Frozen serial 8 μm sections of mice heart tissues were cut at −20°C with a Microtome (CM1950; Leica, Nussloch, Germany) and mounted on ITO-coated glass slides for MALDI-MSI and on coated glass slides for HE staining. The sections were dehydrated in a 50 ml conical tube containing silica gel and stored at −80°C.

The prepared matrix solution was applied over 200 µL/section (9AA) with an airbrush (GSI Creos, Tokyo, Japan). After spraying, the samples were immediately measured.

MALDI-MSI experiment was performed on a MALDI ion trap time-of-flight mass spectrometer (iMScope TRIO; Shimadzu, Kyoto, Japan) equipped with a 1-kHz Nd:YAG laser (λ = 355 nm). The laser spot size was approximately 15 µm, and each pixel was irradiated 80 times at a repetition rate of 1 kHz. For m/z calibration, Polyethylene Glycol 600 Sulfate (Tokyo Chemical Industry, Tokyo, Japan) was used. Mass spectra were acquired in the negative ion detection mode. The target m/z values were 505.99 derived from [M-H]^-^. In the imaging experiment, the interval of data points was 40 µm in the lateral and axial directions. After sample analysis, ion images were reconstructed based on data extracted from m/z ranges of the target m/z values ± 0.02 Da using Imaging MS solution (Shimadzu, Kyoto, Japan). Signal intensities from region-of-interest were extracted using Imaging MS solution.

### Mathematical Model

We constructed the mathematical model of TNFR signaling pathway by extending the previous model ^67^. A comprehensive diagram of the TNFα signaling network is shown in Figure. S3a. The mathematical model describes the biochemical reactions using ordinary differential equations, from activation of TNF receptors following transcription of negative feedback regulators (IκBα and A20) by NFκB. The model constitutes 17 ordinary differential equations with 34 parameters and 18 model species. Details about the model equations can be found in Supplementary Table 1–3.

Parameter estimation was performed using BioMASS (ver. 0.7.2) ^68^ (https://github.com/biomass-dev/biomass) by minimizing the sum of cosine similarity between experimental data and simulated values in the genetic algorithm (https://ui.adsabs.harvard.edu/abs/2003TJSAI..18..193K/abstract).

The cosine similarity between 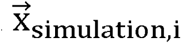 and 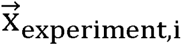, is defined as

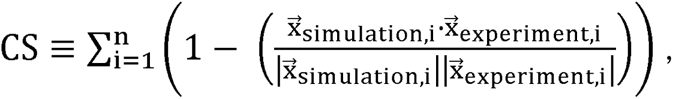

where CS is cosine similarity, n = 3 is the number of molecules used in parameter estimation (IKK, I B, and NF B), 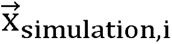 is a vector for the activity of the molecules at each time point in the simulation, 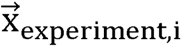 is vector for activities of the molecule at each experimental time point. 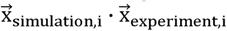 is the inner product of 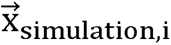 and 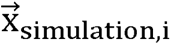, and |·| denotes the magnitude of a vector. A low cosine similarity means that the dynamic behaviors of the molecules in simulation are similar to the ones in the experiment. We used 30 parameter sets for our model simulation and analysis.

### Sensitivity analysis

The sensitivity coefficient of each reaction parameter, S_i_, is defined as

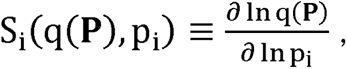

where p_i_ is an i-th parameter, **P** is parameter vector **P** = (p_1_, p_2_,…), and q(**P**) is a target function, for example, sum of signals, and oscillatory amplitude and period. The sum of amplitudes of peaks, and average of periods from the first to the second were also used as target functions for the analysis. The sensitivity was calculated by increasing the parameter values by 1%.

### RNA-sequencing (RNA-seq) of MCF7 cells and mouse heart tissues

Total RNA was isolated using NucleoSpin RNA Plus (Macherey-Nagel, Duren, Germany, Cat. No. 740984.250). Library construction and RNALseq were performed using the Illumina sequencing platform (Rhelixa). Poly(A) RNA was prepared with the Poly(A) mRNA Magnetic Isolation Module (New England Biolabs, Ipswich, MA, USA), the sequencing library was generated with the NEBNext® Ultra™II Directional RNA Library Prep Kit (New England Biolabs) and paired-end sequencing was processed with the NovaSeq 6000 (Illumina, San Diego, CA).

### Acquisition of time course RNA-seq data

Time course bulk RNA-seq data sets (DRA011742 and DRA011743 of DNA Data Bank of Japan) in which gene expression was measured every 15 min after TNFα stimulation in IκBα KD MCF-7 cells were obtained from our previous study ^19^.

### MCF-7 RNA-seq data analysis

For analysis of MCF7 RNA-seq, sequencing reads were preprocessed using nf-core/rnaseq pipeline (v.3.5) ^69^ (https://github.com/nf-core/rnaseq/blob/1.4.2/docs/output.md). Read quality was assessed using FastQC (v.0.11.9). Trim Galore (v.0.6.7) was used for adaptor trimming. The resulting reads were mapped onto the GRCh38 genome using STAR (v.2.7.6a) and then quantified by Salmon (v.1.5.2). Protein coding genes registered in GENCODE (https://www.gencodegenes.org/human/release_40.html) were used for downstream analysis. Differentially Expressed Genes (DEGs) were identified using the R package DESeq2 ^70^ by performing a Wald significance test between gene expression levels in each condition 48 h after TNFα stimulation and calculating adjusted P-values and log_2_ fold change (TNFα vs IκBαKD+TNFα). Genes with adjusted P-value < 0.01 and log_2_ fold change > 0.4 were classified as upregulated DEGs, and genes with adjusted P-value < 0.01 and log_2_ fold change < −0.4 as downregulated DEGs. The expression pattern of each upregulated DEG in Figure 3B was obtained using time course TPM data. Time course TPM data were Z-scored after the logarithm of 2 was taken before clustering. Clustering was performed using the partitioning around medoids (PAM) algorithm with the R package cluster. Heat maps were created using the R package complexHeatmap ^71^. The creation of the regression line and the calculation of the 95% confidence intervals were calculated by the function “stat_smooth()” of the R package ggplot2 ^72^.

### Mouse heart RNA-seq data analysis

Mapping of data to perform Gene/transcript quantification was performed using nf-core/rnaseq pipeline (v3.5.) and the default GRCm38 genome. For TF activity analysis, the DOROTHEA ^73^ Regulon database was used in combination with VIPER ^74^. Only TFs with a confidence score of A and interacting with at least 30 genes were included. Genes that were significantly upregulated with aging were identified using the R package DESeq2 ^70^. Criterion was set as adjusted P-values < 0.01 and log_2_ fold change > 0.

### Assay for transposase-accessible chromatin with high-throughput sequencing (ATAC-seq)

Library preparation, sequencing, and mapping for ATAC-seq were performed at DNAFORM (Yokohama, Japan). Fragmentation and amplification of ATAC-seq libraries were conducted as described previously ^75^. Briefly, approximately 5×10^4^ cells were lysed and transposed reactions were carried out using Tn5 Transposase (Illumina, #FC121-1030) at 37 °C for 30 min. The reaction solution was purified with the Qiagen MinElute PCR Purification Kit. PCR with custom Nextera PCR primers was then performed for five cycles using NEBNext Q5 Hot Start HiFi PCR Master Mix (New England Biolabs, Ipswich, MA, USA) ^76^. The number of additional PCR cycles was determined by qPCR of partially amplified products ^76^. The PCR products were purified using Agencourt AMPure XP beads (Beckman Coulter: A63881) by double size selection (left ratio: 1.4x, right ratio: 0.5x) following the manufacturer’s protocol. Paired-end sequencing was performed on the Illumina HiSeq sequencer.

### MCF7 ATAC-seq data analysis

Mapping and peak calls were conducted by ENCODE ATAC-seq pipeline (https://github.com/ENCODE-DCC/atac-seq-pipeline). Reads were mapped to the GRCh38 reference sequence using Bowtie2 (v.2.3.4.3), and duplicate reads were removed with Picard (v.2.20.7) and samtools (v.1.9). Peak calling was performed using MACS2 (v.2.2.4) with default parameters. After removal of the blacklist region, the consistency of peaks was tested by Irreproducible Discovery Rate (IDR) using IDR (v.2.0.4.2). Peak annotation was conducted by HOMER (v.4.9.1) with default settings. Changes in chromatin accessibility near the transcription start site (TSS ± 5000 bp) per condition in each region and the list of differential accessible peaks that were differentially accessible under IκBα KD+TNFα conditions compared to controls were obtained using DEseq2 (v.1.20.0). The criteria for determining a peak with a difference in accessibility were adjusted P-value < 0.05 and log_2_ fold change > 0.

### RELA chromatin immunoprecipitation-sequencing (ChIP-seq)

ChIP was carried out using the Simple ChIP Enzymatic Chromatin IP Kit (Cell Signaling Technology, #9005) following the manufacturer’s protocol. Protein/DNA was crosslinked using 1% formaldehyde. Nuclei were isolated and sheared using an ultrasonicator (Epishear probe sonicator, Active Motif, Carlsbad, CA, USA). The sheared chromatin was incubated with RELA antibody (Abcam, ab7970) and Dynabeads Protein G beads (10004D, Thermo Fisher Scientific). After elution of ChIP products, DNA was purified using MinElute PCR Purification Kit (QIAGEN, 28004), the library was prepared using the TruSeq ChIP Library Preparation Kit (Illumina), and 36-bp single-end sequencing of the libraries was processed with a HiSeq3000 system (Illumina).

### RELA ChIP-seq data analysis

Mapping of data to produce BigWig files and call peaks was performed using the nf-core/chipseq pipeline (v.1.2.2) ^69^ (https://doi.org/10.5281/zenodo.3240506) and default GRCh38 genome. BigWig files were quantified by “computeMatrix” and visualized by “plotHeatmap” and “plotPlofile” in deeptools (v3.5.1) ^77^. Each peak was assigned to the TSS of the closest gene by HOMER (v.4.11) ^78^. The RELA target gene was defined as the gene whose peak was within ± 5,000 bp from TSS and They were extracted from each cluster. The number of RELA peaks annotated to each target gene was counted using R.

### RELA binding site analysis using public databases

The ChIP Atlas peak browser ^79^ was used to identify RELA binding regions in the mouse genome (mm10). The ChIP-seq peak significance threshold was set to 500 (q value < 1E-50) and ChIP-seq peaks from the groups “TFs and other” and “All cell types” and “RELA” were used. Multiple RELA ChIP peaks in a single region were merged by mergeBed in deeptools (v.3.5.1)^80^. These RELA ChIP peaks were then assigned to the TSS of the closest gene by HOMER (v.4.11). The final RELA target gene was defined as the gene whose Rela peak was within 500 bp of the TSS. The number of RELA peaks within ±10,000 bp from the TSS annotated to each target gene was then counted on R.

### KEGG enrichment analysis

KEGG enrichment was compared by the “comparecluster()” function in clusterProfiler ^81^ R package with “fun = “enrichKEGG”. The result was plotted by the R package “complexHeatmap”. Terms with adjusted P-values less than 0.05 were displayed.

### Data and materials availability

Metabolome data are included in the Supplementary Data. The sequence data for RNA-seq, ChIP-seq, and ATAC-seq reported in this paper have been deposited in the DNA Data Bank of Japan with accession number DRA015837. Code is available at https://github.com/okadalabipr. All other data will be available upon reasonable request.

## QUANTIFICATION AND STATISTICAL ANALYSIS

We performed all experiments at least twice and confirmed similar results. Statistical analyses were performed using GraphPad Prism ver. 8.0 software (GraphPad Software, Inc., La Jolla, CA, USA) and R studio ver. 4.2.2 (Rstudio, Boston, MA, USA). For *in vitro* experiments, data from two or more groups were analyzed using Student’s t-tests and one-way analysis of variance (ANOVA), respectively. For *in vivo* experiments, data from two or more groups were analyzed using the Mann–Whitney U and Kruskal–Wallis tests, respectively. Data represented as mean ± SEM or ± SD; P values < 0.05 were considered statistically significant.

## Supplementary Figures

**Figure S1. Effect of IκBα knockdown/knockout on cellular senescence (related to Figure 1).** (A) IκBα knockdown in MCF-7 cells by siRNA. MCF-7 cells were treated with TNFα (10 or 30 ng/mL) for 24 or 48 hr. Representative blot images are shown. The values under each lane indicate the relative density of the band normalized to GAPDH. (B) Representative images of RELA immunofluorescence staining in MCF-7 cells transfected with control and IκBα siRNA. The cells were treated with 10 ng/mL TNFα for the indicated times. Scale bars, 25 μm. (C) Principal component analysis of 13 geometric parameters for single cells (D) Changes of the geometric parameters for single cells. (E) IκBα knockout (KO) in MCF-7 cells by CRISPR-Cas9. (F) Effect of IκBα KO on cell viability in MCF-7 cells treated with TNFα (10 ng/mL) for 48 hr. (G) Effect of IκBα KO on SA-β-gal staining in MCF-7 cells treated with TNFα for 24 hr. (H) Knockdown of RELA and IκBα in MCF-7 cells by siRNA. MCF-7 cells were treated with TNFα (10 ng/mL) for 48 hr. (I and J) Cell cycle analysis of MCF-7 cells under IκBα KD+TNFα conditions. MCF-7 cells transfected with the control and IκBα siRNA and treated with 10 ng/mL TNFα for the indicated times. Representative images of FUCCI immunofluorescence (I). MCF-7 cells show G1-phase (green color; PIP-tag-mNeonGreen), S-phase (magenta color; mScarlet-hGem) and G2/M-phase (merged color; PIP-tag-mNeonGreen and mScarlet-hGem). Scale bars, 100 μm. Cell cycle dynamics mapped at an individual cell level (N = 24) (J). Cell cycle phase data were obtained for cells that could be continuously tracked for 60 hr after TNFα treatment. As the experiment was performed under non-synchronized cell cycle conditions, an equal number of cells starting from the G1 or S/G2/M phase (N = 12 for each) were selected for analysis. Data represent the mean ± SD [(F) and (G)]. *P* values were determined using one-way ANOVA followed by Tukey’s test [(D), (F), and (G)].

**Figure S2. Model simulation of NFκB dynamics under knockdown or overexpression of IκBα or A20 (related to Figure 2).** (A) Effect of A20 knockdown (KD) on protein expression of A20 and Cyclin D1 in MCF-7 cells. Representative blot images are shown. Asterisks indicate a non-specific band. (h) Effect of A20 KD on SA-β-gal staining in MCF-7 cells treated with TNFα (10 ng/mL). Representative images are shown. Scale bars, 100 μm. (B) Detailed diagram of the TNFα signaling network. Each box represents a regulatory module leading from the IKK-IκBα-NFκB module to the transcription and translation module. Black arrows represent biochemical modifications, association/dissociation, and transport of proteins. Red numbers indicate model reaction indices. (C) Protein expression of nuclear RELA in MCF-7 cells treated with TNFα (10 ng/mL) for the indicated time (upper panel). (D) Protein expression of P-IKKα/β, IKKβ, and IκBα in MCF-7 cells treated with TNFα for the indicated time (lower panel). (E) Time-course patterns of P-IKK, total IκBα, and total nuclear NFκB in MCF-7 cells treated with TNFα (10 ng/mL) for the indicated time (upper). Results of model predictions are shown (IκBα and A20 mRNA) (lower). Circles represent experimental data (mean ± SD, N = 3) and lines represent the averaged trajectories of 30 simulation runs. The darker colored line represents the average of the simulation results using 30 independently searched parameter sets. The lighter-colored lines represent the simulation results for the individual parameter sets. (F) Estimated 30 parameter sets. The box plot shows the range of values for 30 parameter sets estimated from the time-course experimental datasets shown in Figure S2D. The horizontal axis shows the indices of the estimated model parameters. The vertical axis shows the parameter values and protein abundances (TTR, IKK, and IκBα-NFκB). (G) Simulations of nuclear NFκB abundance in IκBα- or A20-overexpressing cells treated with TNFα for the indicated time. The color bar indicates the fold change values of the IκBα or A20 transcription rate compared to that of the control. The data were normalized between the minimum (0) and maximum (1) values in the control. (H) Sensitivity analysis against nuclear NFκB activity. Nuclear NFκB activity was defined as the area under the curve (AUC), which represents the sum of the NFκB signals (left). Sensitivity of each parameter to nuclear NFκB activity (right). Larger changes in the graph indicate larger sensitivities against the 1% change in reaction parameters in the model diagram in Figure S2B. Error bars indicate the mean±SD of the 30 parameter sets. (I) Model simulation of the NFκB dynamics under IκBα KD and A20 overexpression conditions. Simulations of nuclear NFκB dynamics under IκBα KD and A20 overexpression conditions in response to TNFα. Each line is presented as the mean of simulation values for 30 parameter sets. The color bar indicates the fold change values of the IκBα transcription rate compared to that of the control. The data were normalized between the minimum (0) and maximum (1) values in the control. (J) Overexpression of A20 in MCF-7 cells. Representative blot image of A20 and EGFP protein. Asterisks indicate a non-specific band. (K) Effect of IκBα KD and TNFα on the time-course nuclear RELA abundance in A20 OE MCF-7 cells.

**Figure S3. MA plot of the control vs IκBα KD+TNFα ATAC-seq signals (related to Figure 3).** Each dot represents each open chromatin region, and red dots indicate open chromatin regions identified by padj < 0.05 and log_2_ FC > 0.

**Figure S4. Expression of IκBα, A20, and TNFα in young and aged mouse tissues. (related to Figure 4).** (A–D) Protein expression of IκBα, A20, and TNFα in young and aged mouse tissues. Young mice: 8 weeks old, N = 8. Aged mice: 69–73 weeks old, N = 8. Representative blot images are shown (A). Box plots quantifying the relative protein level of IκBα (B), A20 (C), and TNFα (D). Data represent the median and interquartile range (25th and 75th percentiles) with error bars showing minimum to maximum. *P* values were determined using the Mann–Whitney U test. (E) Immunohistochemistry of RELA in heart tissues from young and aged mice. Young mice: 8 weeks old, N = 3. Aged mice: 69–73 weeks old, N = 3. Representative images are shown. Scale bars, 50 μm.

**Figure S5. Effect of the AKT inhibitor on Cyclin D1 expression in MCF-7 cells under IκBα+TNFα conditions (related to Figure 5).** Expression of CyclinD1 in IκBα KD MCF-7 cells treated with TNFα and inhibitor. Representative blot images are shown. Cells were treated with BAY11-7085 (5 µM, NFκB inhibitor), AKTi VIII (1 µM, AKT1/2 inhibitor), LY294002 (10 µM, PI3K inhibitor), rapamycin (100 nM, mTOR inhibitor), U01268 (10 µM, MEK inhibitor), SB216763 (10 µM, GSK3β inhibitor), SB203580 (10 µM, inhibitor for p38 and AKT), and TNFα for 48 hr. MG132 (10 μM, proteosome inhibitor) was added 8 hr before collecting samples.

**Figure S6. Effect of sustained NFκB activation on purine metabolism in MCF-7 cells (related to Figure 6).** (A) Principal component analysis of metabolome data (intracellular) of IκBα knockdown (KD) MCF-7 cells treated with TNFα for 48 hr. (B) Volcano plots showing differences in extracellular metabolites in MCF-7 cells under control and IκBα KD conditions. (C) Extracellular hypoxanthine levels in IκBα KD MCF-7 cells treated with TNFα. (D) Effect of IκBα KD on the mRNA expression of purine metabolism genes in MCF-7 cells. The cells were treated with TNFα (10 ng/mL) for 48h. (E) HPRT1 KD in MCF-7 cells by siRNA. Left, representative blot image. Right, bar plots quantifying the relative protein level of HPRT1. Data represent the mean ± SD.

**Figure S7. Protein expression of RICTOR, P-RICTOR, and HPRT1 in young and aged hearts (related to Figure 7).** (A) Protein expression of RICTOR, P-RICTOR, and HPRT1 in young and aged mouse tissues. Young mice: 8 weeks old, N = 8. Aged mice: 69–73 weeks old, N = 8. Representative blot images are shown. (B) Metabolite pathway enrichment analysis for the differential metabolites between the heart tissues of young and aged mice. (C) Mass spectrum of the mouse heart. Negative ion mode (m/z 340 to m/z 530) mass spectrum of the mouse heart tissue. Asterisks indicate the ATP peak. (D) Effect of H_2_O_2_ on IκBα protein expression in MCF-7 cells. MCF-7 cells were treated with H_2_O_2_ for 96 hr. The values under each lane indicated the relative density of the band normalized to ERK.

