## Supplementary figures for "Nuclear NFκB Activity Balances Purine Metabolism in Cellular Senescence"

### Supplementary figure 1

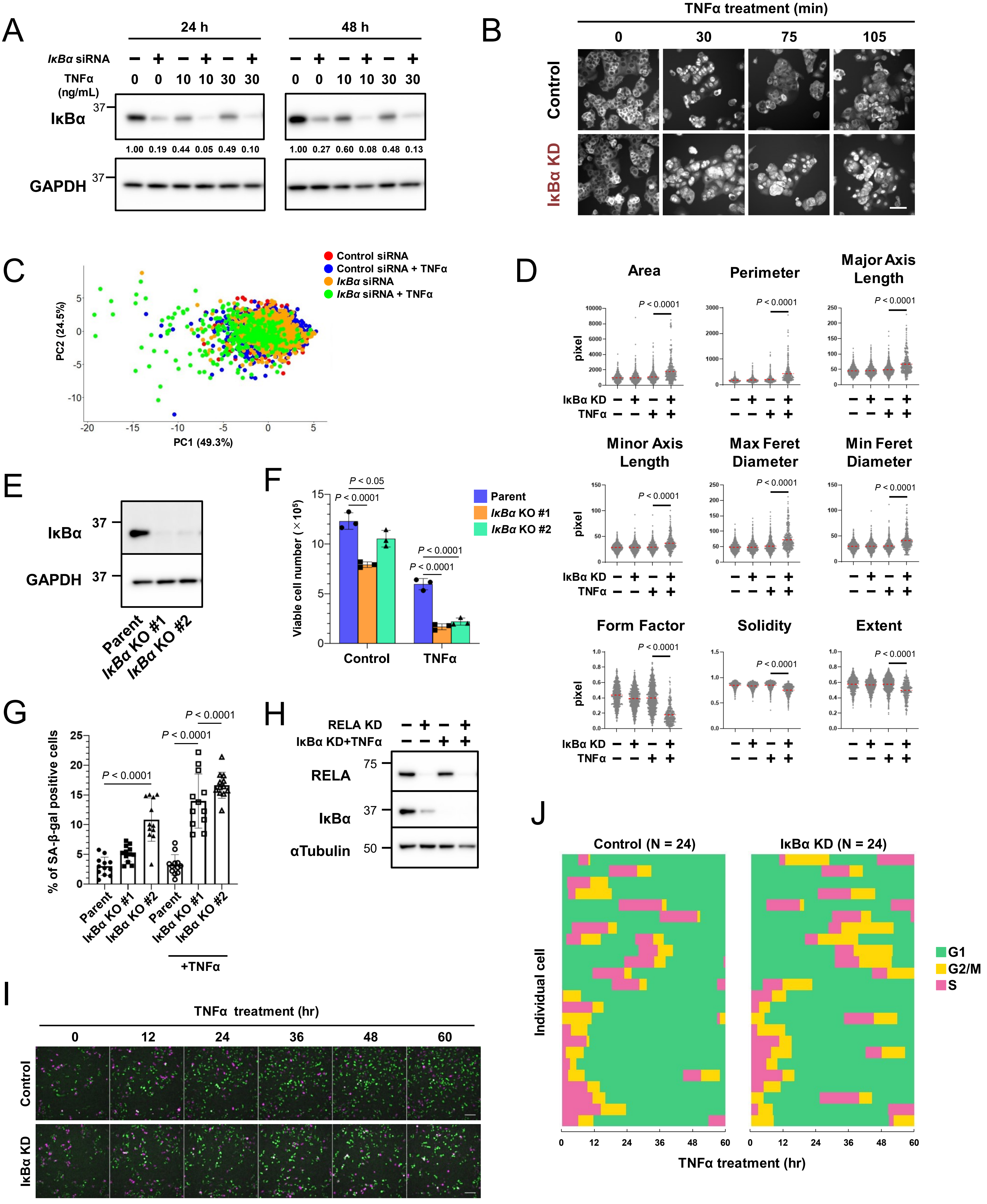

### Supplementary figure 2

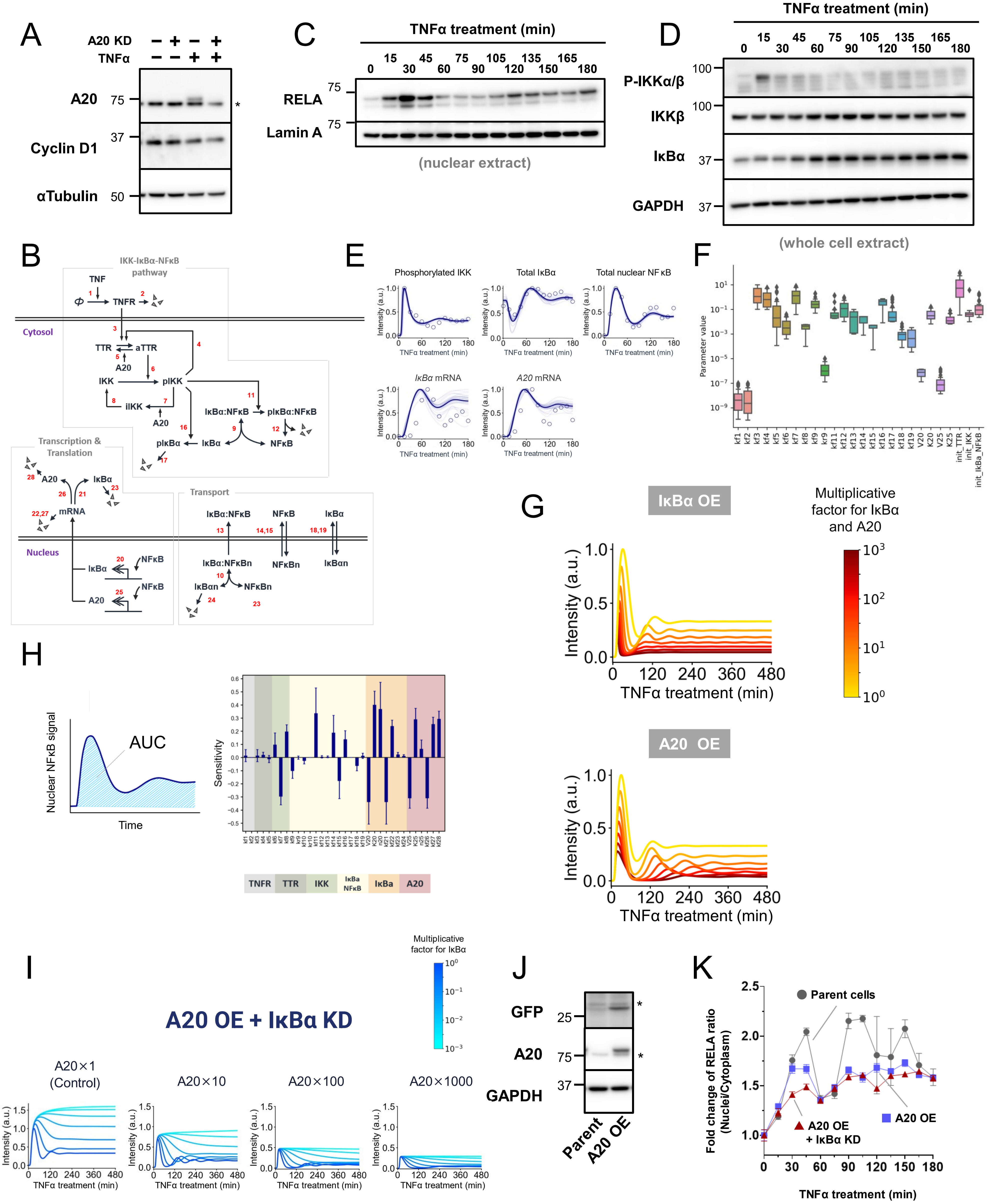

### Supplementary figure 3

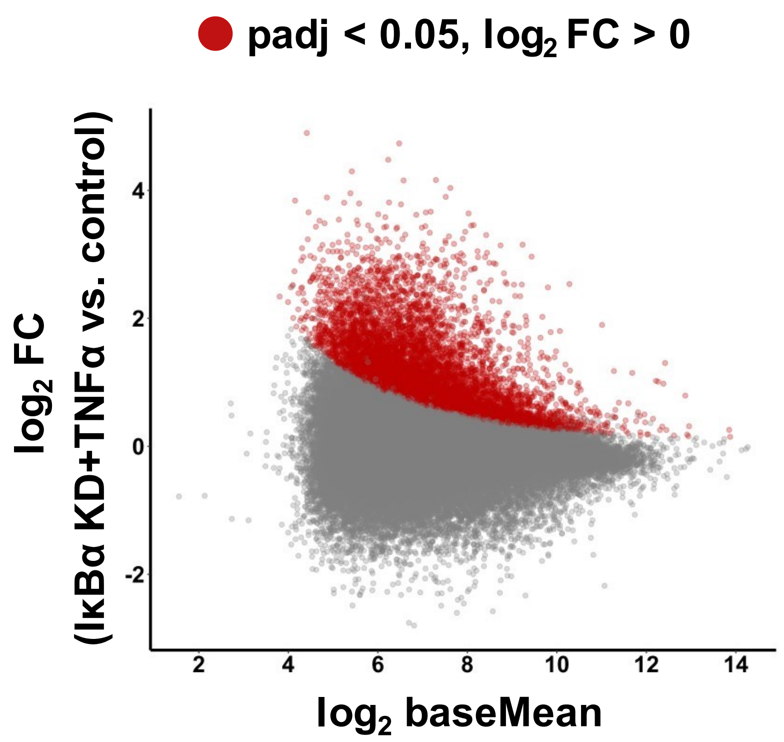

### Supplementary figure 4

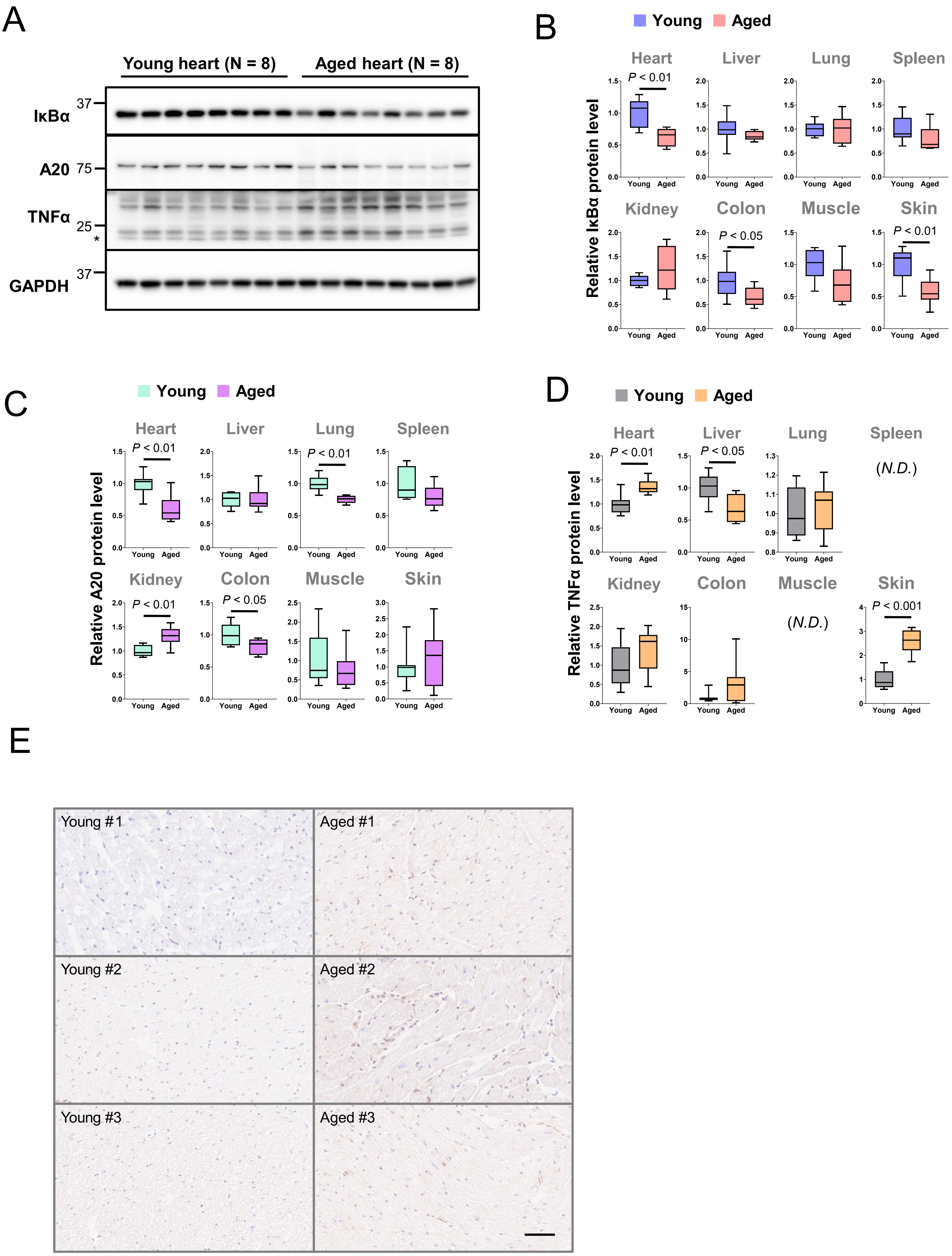

### Supplementary figure 5

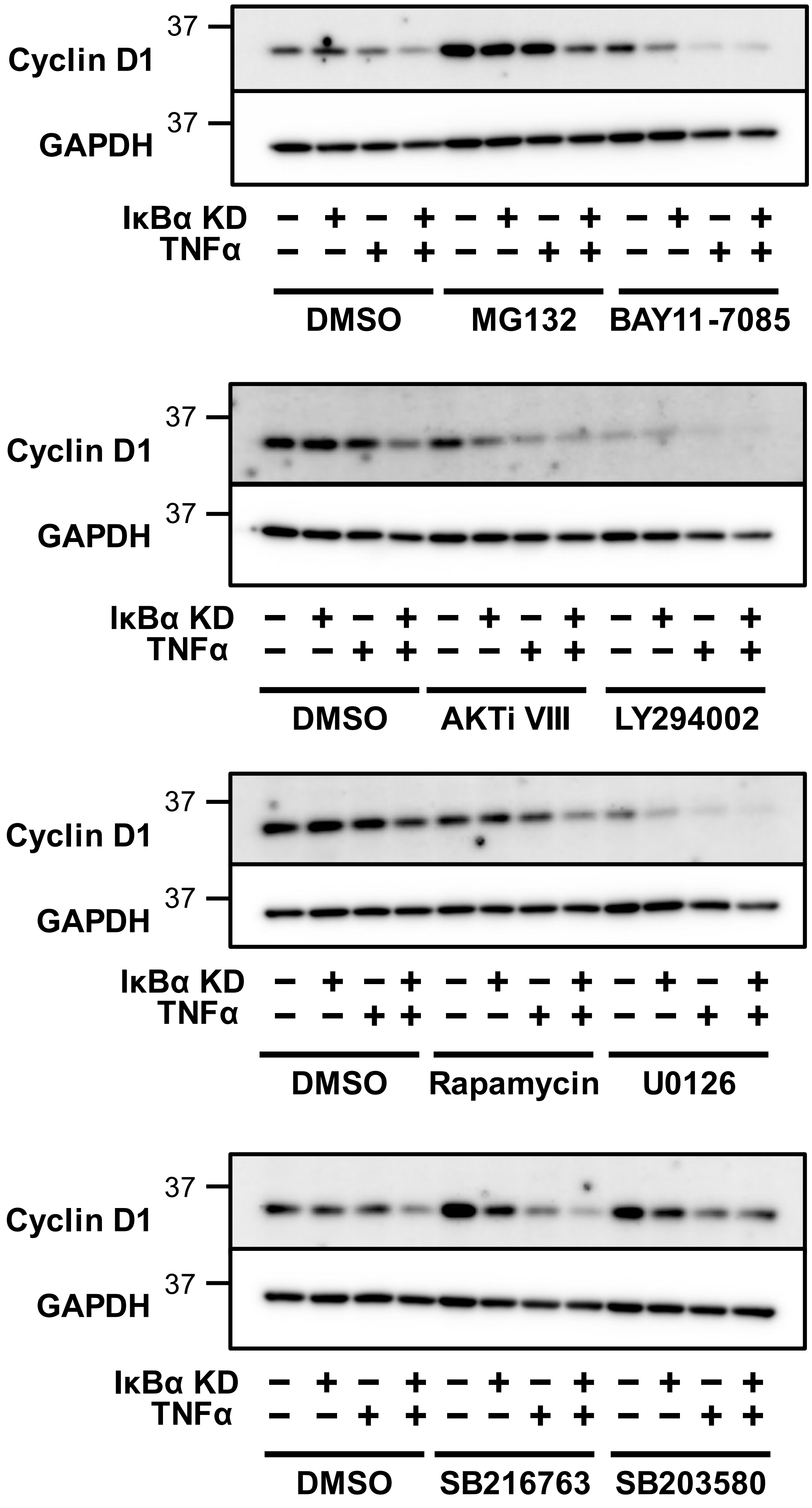

### Supplementary figure 6

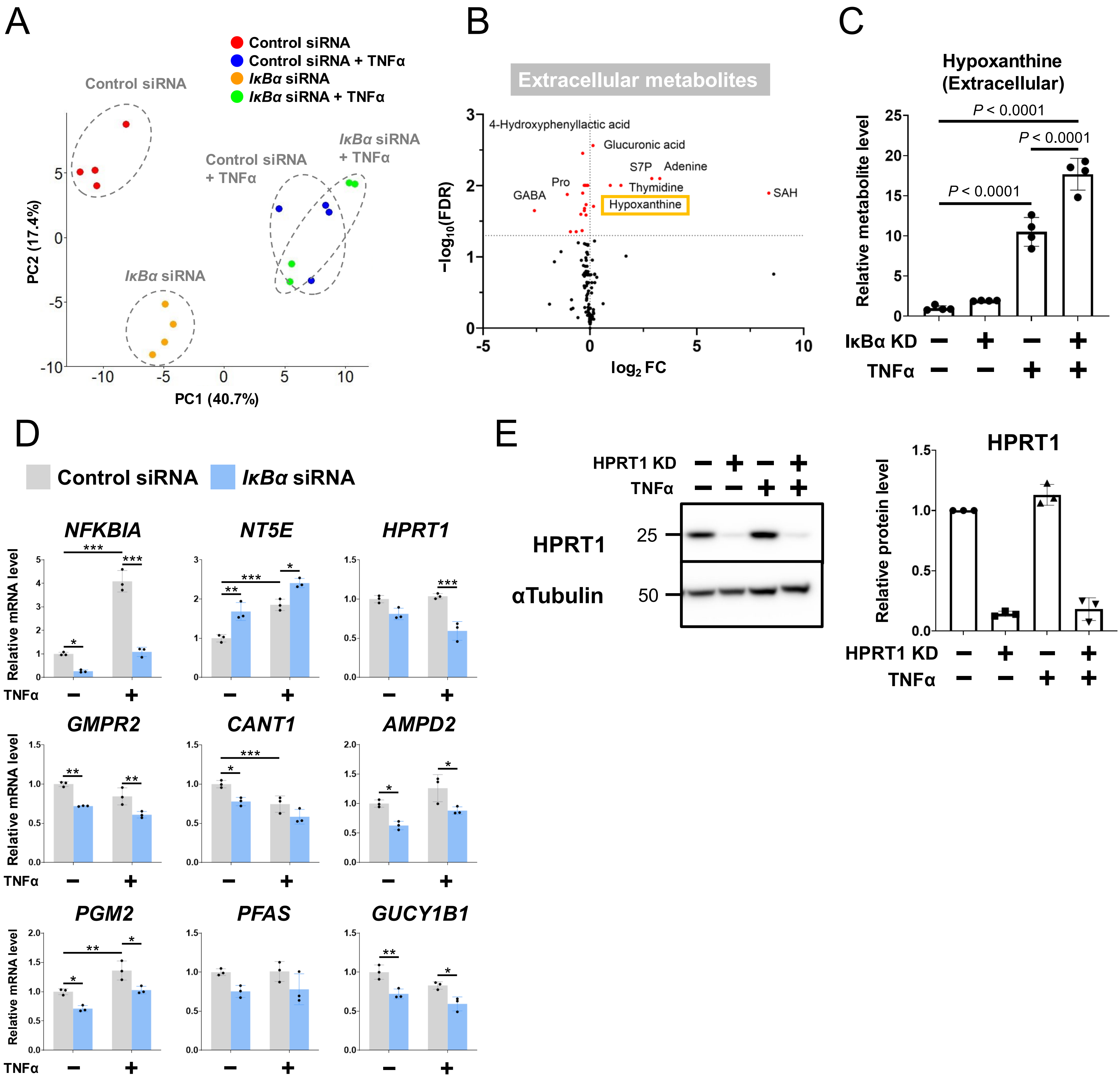

### Supplementary figure 7

A

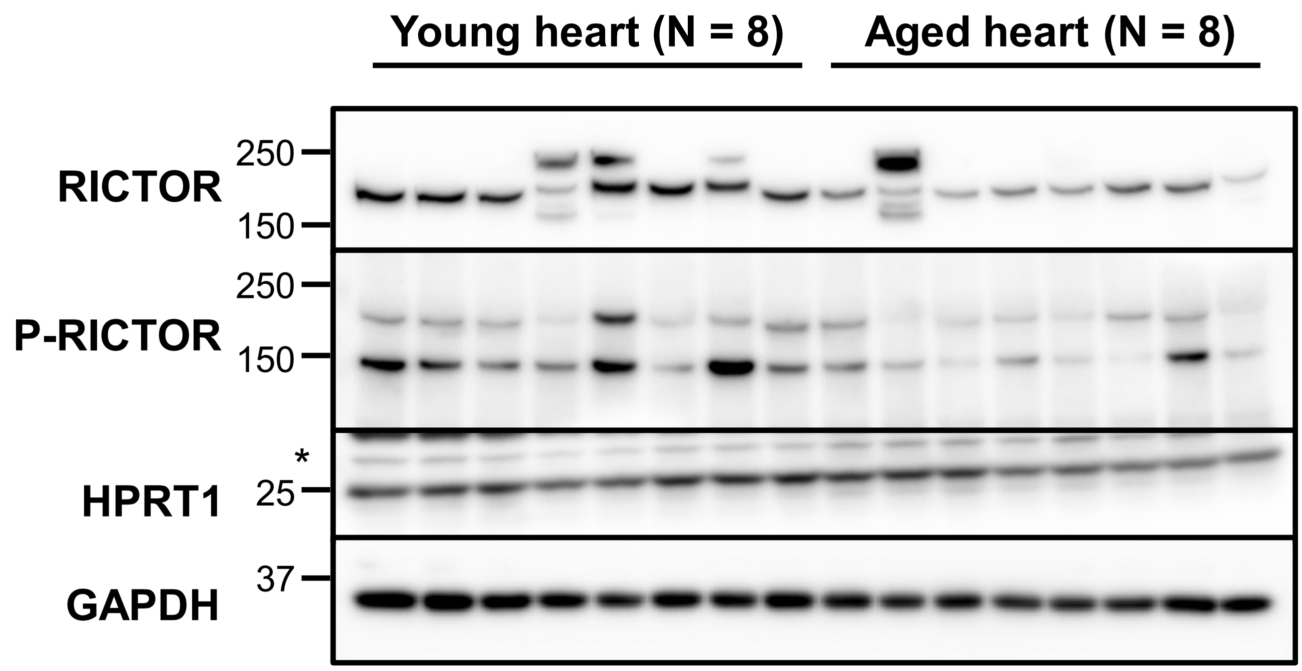


B

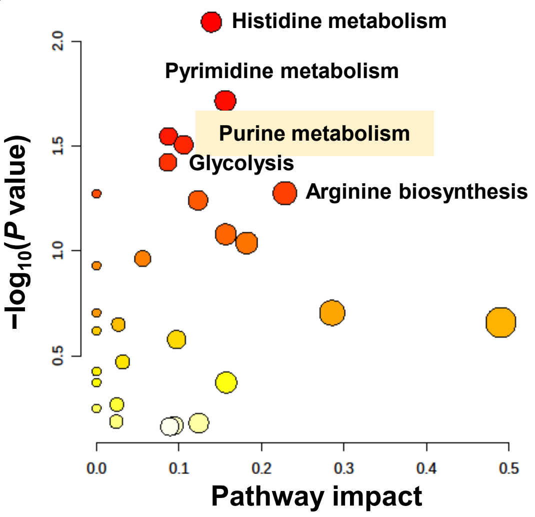


C

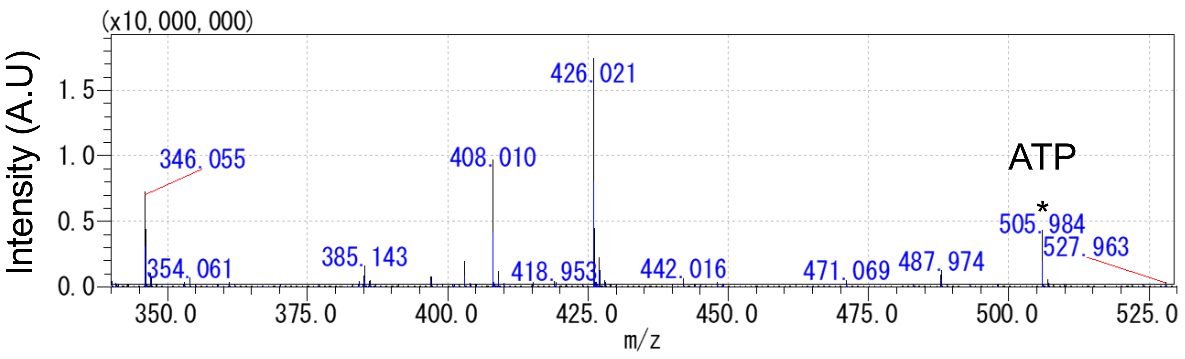


D

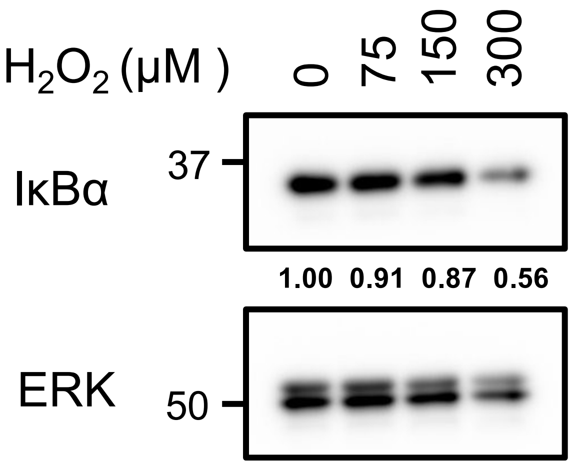
