## Supplementary Tables. for "Nuclear NFκB Activity Balances Purine Metabolism in Cellular Senescence"

Supplementary Table 1. Reactions.

|  |  |  |  | Rate |
| --- | --- | --- | --- | --- |
| $v1$ | ∅ | → | TNFR | $k1･[TNF]$ |
| $v2$ | TNFR | → | ∅ | $k2･[TNFR]$ |
| $v3$ | TTR | → | aTTR | $k3･\left[ TNFR \right]･[TTR]$ |
| $v4$ | TTR | → | aTTR | $k4･\left[ pIKK \right]･[TTR]$ |
| $v5$ | aTTR | → | TTR | $k5･\left[ A20 \right]･[aTTR]$ |
| $v6$ | IKK | → | pIKK | $k6･\left[ aTTR \right]･[IKK]$ |
| $v7$ | pIKK | → | iIKK | $k7･\left[ A20 \right]･[pIKK]$ |
| $v8$ | iIKK | → | IKK | $k8･\left[ iIKK \right]$ |
| $v9$ | IκBα+NFκB | → | IκBα:NFκB | $kf9･\left[ IkBa \right]･\left[ NFkB \right]-kr9･\left[ IkBa:NFkB \right]$ |
| $v10$ | IκBαn+NFκBn | → | IκBα:NFκBn | $kf10･\left[ IkBan \right]･\left[ NFkBn \right]-kr10･\left[ IkBa:NFkB \right]$ |
| $v11$ | IκBα:NFκB | → | pIκBα:NFκB | $k11･\left[ pIKK \right]･\left[ IkBa:NFkB \right]$ |
| $v12$ | pIκBα:NFκB | → | NFκB | $k12･\left[ pIkBa:NFkB \right]$ |
| $v13$ | IκBα:NFκBn | → | IκBα:NFκB | $k13･[IkBa:NFkBn]$ |
| $v14$ | NFκB | → | NFκBn | $k14･[NFkB]$ |
| $v15$ | NFκBn | → | NFκB | $k15･[NFkBn]$ |
| $v16$ | IκBα | → | pIκBα | $k16･\left[ pIKK \right]･\left[ IkBa \right]$ |
| $v17$ | pIκBα | → | ∅ | $k17･\left[ pIkBa \right]$ |
| $v18$ | IκBα | → | IκBαn | $k18･[IkBa]$ |
| $v19$ | IκBαn | → | IκBα | $k19･[IkBan]$ |
| $v20$ | ∅ | → | *IκBα* mRNA | $\frac{{V20･\left[ NFkBn \right]}^{n20}}{{K20}^{n20}+\left[ NFkBn \right]^{n20}}$ |
| $v21$ | *IκBα* mRNA | → | IκBα | $k21･\left[ ikba\_mRNA \right]$ |
| $v22$ | *IκBα* mRNA | → | ∅ | $k22･\left[ ikba\_mRNA \right]$ |
| $v23$ | IκBα | → | ∅ | $k23･\left[ IkBa \right]$ |
| $v24$ | IκBαn | → | ∅ | $k24･\left[ IkBan \right]$ |
| $v25$ | ∅ | → | *A20* mRNA | $\frac{{V25･\left[ NFkBn \right]}^{n25}}{{K25}^{n25}+\left[ NFkBn \right]^{n25}}$ |
| $v26$ | *A20* mRNA | → | A20 | $k26･\left[ a20\_mRNA \right]$ |
| $v27$ | *A20* mRNA | → | ∅ | $k27･\left[ a20\_mRNA \right]$ |
| $v28$ | A20 | → | ∅ | $k28･\left[ A20 \right]$ |

Supplementary Table 2. List of equations of the model

Equations

$$\frac{d\left[ TNFR \right]}{dt}=+v1-v2$$

$$\frac{d[TTR]}{dt}=-v3-v4+v5$$

$$\frac{d[aTTR]}{dt}=+v3+v4-v5$$

$$\frac{d\left[ IKK \right]}{dt}=-v6+v8$$

$$\frac{d\left[ pIKK \right]}{dt}=+v6-v7$$

$$\frac{d\left[ iIKK \right]}{dt}=+v7-v8$$

$$\frac{d[NFkB]}{dt}=-v9+v12-v14+v15$$

$$\frac{d\left[ NFkBn \right]}{dt}=-v10+v14+v15$$

$$\frac{d[IkBa]}{dt}=-v9-v16-v18+v19+v21-v23$$

$$\frac{d[IkBan]}{dt}=-v10+v18-v19-v24$$

$$\frac{d[IkBa:NFkB]}{dt}=+v9-v11+v13$$

$$\frac{d\left[ IkBa:NFkBn \right]}{dt}=+v10-v13$$

$$\frac{d[pIkBa]}{dt}=+v16-v17$$

$$\frac{d\left[ pIkBa:NFkB \right]}{dt}=+v11-v12$$

$$\frac{d\left[ ikba\_mRNA \right]}{dt}=+v20-v22$$

$$\frac{d[a20\_mRNA]}{dt}=+v25-v27$$

$$\frac{d[A20]}{dt}=+v26-v28$$

Supplementary Table 3. List of observables

| Experiment | Simulation |
| --- | --- |
| Phosphorylated IKK | $\left[ pIKK \right]$ |
| Total IκBα | $\left[ IkBa \right]+ \left[ IkBa:NFkB \right]+ \left[ pIkBa \right]+ \left[ pIkBa:NFkB \right]+ \left[ IkBan \right]+\left[ IkBa:NFkBn \right]$ |
| Total nuclear NFκB | $\left[ NFkBn \right]+\left[ IkBa:NFkBn \right]$ |
| Nuclear NFκB | $\left[ NFkBn \right]$ |
| *IκBα* mRNA | $\left[ ikba\_mRNA \right]$ |
| *A20* mRNA | $\left[ a20\_mRNA \right]$ |

Supplementary Table 4.

Primer sequences.

| Gene | Forward primer sequence (5'-3') | Reverse primer sequence (5'-3') |
| --- | --- | --- |
| *IL8* | CCTGATTTCTGCAGCTCTGTGT | GGTGGAAAGGTTTGGAGTATGTCT |
| *CCL2* | AAGACCATTGTGGCCAAGGA | TTCGGAGTTTGGGTTTGCT |
| *NFKBIA* (*IκBα*) | AGACGAGGAGTACGAGCAGAT | GCCAAGTGCAGGAACGAGT |
| *TNFAIP3* (*A20*) | GAGAGCACAATGGCTGAACA | CACAAGCTTCCGGACTTCTC |
| *NT5E* | GGCACTATCTGGTTCACCGT | CCTCTTTGAGGAGTGGCTCG |
| *HPRT1* | TTGCTTTCCTTGGTCAGGCA | ATCCAACACTTCGTGGGGTC |
| *GMPR2* | GGAGCTGACTTCGTGATGCT | CTCTCGATGAGCTCACCACC |
| *CANT1* | CTGGGTGATTCTGTCCGACG | CGGGTTCTCGTTCACCACAT |
| *AMPD2* | GAGCTGCGGCTCTCCATTTA | CGGTACACATCAAAGAGGCG |
| *PGM2* | CCCTTTGTGCCCTTCACAGTA | CGTGAGGAGAAATGATCTGAGC |
| *PFAS* | CGACGATTCGAGGTGCTCT | GACATCTCTGCAGGTGTCCTT |
| *GUCY1B1* | TCACTCAGTGTGGCAATGCT | ACGAACCAGCGAGAAGACAG |
| *RPL27* | CTGTCGTCAATAAGGATGTCT | CTTGTTCTTGCCTGTCTTGT |

**Supplementary Table 5.**

gRNA sequences of *NFKBIA*.

| Name | gRNA sequence | Genomic location |
| --- | --- | --- |
| gRNA_1 | TAACCTCGCGGAAAACACCG | chr14:35403909 (−) |
| gRNA_2 | CTGGACGACCGCCACGACAG | chr14:35404547 (−) |
| gRNA_3 | ACGGCGGCACGGACTGCTGT | chr14:35404730 (+) |
