## Supplementary material for "Nuclear NFκB Activity Balances Purine Metabolism in Cellular Senescence": Key Resources Table

| REAGENT or RESOURCE | SOURCE | IDENTIFIER |
| --- | --- | --- |
| Antibodies | | |
| Mouse monoclonal anti-GAPDH | Sigma-Aldrich | Cat# MAB374; RRID:AB_2107445 |
| Rabbit polyclonal anti-GAPDH | Proteintech | Cat# 10494-1-AP; RRID:AB_2263076 |
| Rabbit polyclonal anti-αTubulin | Abcam | Cat# ab15246; RRID:AB_301787 |
| Mouse monoclonal anti-IκBα | Cell Signaling Technology | Cat# 4814; RRID:AB_390781 |
| Rabbit monoclonal anti-IκBα | Cell Signaling Technology | Cat# 4812; RRID:AB_10694416 |
| Rabbit monoclonal anti-RELA | Cell Signaling Technology | Cat# 8242; RRID:AB_10859369 |
| Rabbit monoclonal anti-A20 | Cell Signaling Technology | Cat# 5630; RRID:AB_10698880 |
| Mouse monoclonal anti-RICTOR | Santa Cruz Biotechnology | Cat# sc-81538; RRID:AB_2179969 |
| Rabbit monoclonal anti-RICTOR | Cell Signaling Technology | Cat# 5379; RRID:AB_10691452 |
| Rabbit monoclonal anti-P-RICTOR | Cell Signaling Technology | Cat# 3806; RRID:AB_10557237 |
| Rabbit monoclonal anti-AKT | Cell Signaling Technology | Cat# 4685; RRID:AB_2225340 |
| Mouse monoclonal anti-P-AKT(Ser473) | Cell Signaling Technology | Cat# 4051; RRID:AB_331158 |
| Rabbit monoclonal anti-P-AKT(Ser473) | Cell Signaling Technology | Cat# 4060; RRID:AB_2315049 |
| Rabbit monoclonal anti-P-AKT(Thr308) | Cell Signaling Technology | Cat# 2965; RRID:AB_2255933 |
| Rabbit monoclonal anti-NT5E | Cell Signaling Technology | Cat# 13160; RRID:AB_2716625 |
| Rabbit polyclonal anti-HPRT1 | Proteintech | Cat# 15059-1-AP; RRID:AB_10638622 |
| Mouse monoclonal anti-HPRT1 | Santa Cruz Biotechnology | Cat# sc-376938; RRID:AB_2938532 |
| Rabbit monoclonal anti-Cyclin D1 | Cell Signaling Technology | Cat# 2978; RRID:AB_2259616 |
| Rabbit monoclonal anti-IKKβ | Cell Signaling Technology | Cat# 2370; RRID:AB_2122154 |
| Rabbit monoclonal anti-P-IKKα/β | Cell Signaling Technology | Cat# 2697; RRID:AB_2079382 |
| Mouse monoclonal anti-Lamin A | Abcam | Cat# ab8980; RRID:AB_306909 |
| Rabbit polyclonal anti-ERK | Cell Signaling Technology | Cat# 9102; RRID:AB_330744 |
| Rabbit monoclonal anti-P-ERK | Cell Signaling Technology | Cat# 4370; RRID:AB_2315112 |
| Goat polyclonal anti-GFP | Santa Cruz Biotechnology | Cat# sc-5385; RRID:AB_641121 |
| Chemicals, Peptides, and Recombinant Proteins | | |
| TNFα | Thermo Fisher Scientific | Cat# PHC3015 |
| MG132 | Merck Millipore | Cat# 474790 |
| Rapamycin | Merck Millipore | Cat# 553210 |
| LY294002 | Merck Millipore | Cat# 440202 |
| BAY11-7085 | Fujifilm Wako Pure Chemical Industries, Ltd. | Cat# 020-14331 |
| AKT inhibitor VIII | Cayman Chemical | Cat# 14870 |
| SB216763 | Sigma-Aldrich | Cat# S3442 |
| SB203580 | Cell Signaling Technology | Cat# 5633 |
| U0126 | MedChemExpress | Cat# HY-12031 |
| APCP | Tocris Bioscience | Cat# 3633 |
| Deposited Data | | |
| MCF7 RNA-seq data | This paper | DDBJ: DRA015837 |
| Mouse heart RNA-seq data | This paper | DDBJ: DRA015837 |
| Time course RNA-seq data | https://www.nature.com/articles/s41540-021-00204-7 | DDBJ: DRA011742 and DRA011743 |
| MCF7 ATAC-seq data | This paper | DDBJ: DRA015837 |
| RELA ChIP-seq data | This paper | DDBJ: DRA015837 |
| Experimental Models: Cell Lines | | |
| Human: MCF7 cells | ATCC | HTB-22 |
| Experimental Models: Organisms/Strains | | |
| Mouse: C57BL/6 N mice | CLEA Japan | N/A |
| Oligonucleotides | | |
| Primers for Real-time PCR, see Table S4 | This paper | N/A |
| Control siRNA | Dharmacon/Horizon Discovery | D-001810-02 |
| *IκBα* siRNA | Dharmacon/Horizon Discovery | ON-TARGET plus SMART pool: L-004765-00 |
| *A20* siRNA | Dharmacon/Horizon Discovery | ON-TARGET plus SMART pool: L-009919-00 |
| *RICTOR* siRNA#1 | Dharmacon/Horizon Discovery | ON-TARGET plus SMART pool: L-016984-00 |
| *HPRT1* siRNA#3 | Dharmacon/Horizon Discovery | ON-TARGET plus SMART pool: L-008735-00 |
| gRNA sequences of NFKBIA, see Table S5 | This paper | N/A |
| Recombinant DNA | | |
| pPBbsr2-H2B-iRFP-P2A-mScarlet-I-hGem-P2A-PIP-tag-NLS-mNeonGreen | This paper | https://benchling.com/s/seq-LPZ1tLdpgnpJIYOG2ujR |
| hyPBase | https://www.nature.com/articles/s41467-021-27458-3 | https://benchling.com/s/seq-oGkw53b41IZqvzF5yQ9K |
| eSpCas9(1.1)-T2A-Puro | Addgene | # 101039 |
| A20 cDNA: pPB[Exp]-EGFP/Puro-CMV>hTNFAIP3 | This paper | N/A |
| Software and Algorithms | | |
| MultiExperiment Viewer (MeV) _4_8 ver.10.2 | https://www.future-science.com/doi/abs/10.2144/03342mt01?url_ver=Z39.88-2003&rfr_id=ori:rid:crossref.org&rfr_dat=cr_pub%20%200pubmed | https://sourceforge.net/projects/mev-tm4/ |
| Microsoft excel | Microsoft Corporation | N/A |
| GraphPad Prism v5.0 software | GraphPad Software Inc. | https://www.graphpad.com/scientific-software/prism/ |
| R | The R Project for Statistical Computing | https://www.r-project.org/ |
| R Studio | RStudio | https://www.rstudio.com/ |
| ImageJ (Fiji) | https://doi.org/10.1038/nmeth.2019 | https://imagej.net/software/fiji |
| CHOPCHOP | https://academic.oup.com/nar/article/47/W1/W171/5491735 | https://chopchop.cbu.uib.no/ |
| CellProfiler (ver. 3. 1. 9) | https://journals.plos.org/plosbiology/article?id=10.1371/journal.pbio.2005970 | https://cellprofiler.org/ |
| NanozoomerS210 | Hamamatsu Photonics | https://www.hamamatsu.com/jp/ja.html |
| MetaboAnalyst 5.0 | https://academic.oup.com/nar/article/49/W1/W388/6279832?login=true | https://www.metaboanalyst.ca/ |
| biomaRt (ver. 2.50.3) | https://doi.org/10.1038/nprot.2009.97 | https://bioconductor.org/packages/release/bioc/html/biomaRt.html |
| clusterProfiler (ver. 4.2.2) | https://doi.org/10.1089/omi.2011.0118 | https://bioconductor.org/packages/release/bioc/html/clusterProfiler.html |
| ComplexHeatmap (ver2.10.0) | https://doi.org/10.1093/bioinformatics/btw313 | https://bioconductor.org/packages/release/bioc/html/ComplexHeatmap.html |
| DESeq2 (ver 1.34.0) | https://doi.org/10.1186/s13059-014-0550-8 | https://bioconductor.org/packages/release/bioc/html/DESeq2.html |
| rtracklayer (ver 1.54.0) | https://doi.org/10.1093/bioinformatics/btp328 | https://bioconductor.org/packages/release/bioc/html/rtracklayer.html |
| dorothea (ver 1.6.0) | https://doi.org/10.1101/gr.240663.118 | https://bioconductor.org/packages/release/data/experiment/html/dorothea.html |
| nf-core/rnaseq (ver 3.5) | https://doi.org/10.1038/s41587-020-0439-x | https://github.com/nf-core/rnaseq |
| nf-core/chipseq (ver 1.2.2) | https://doi.org/10.1038/s41587-020-0439-x | https://github.com/nf-core/chipseq |
| ENCODE ATAC-seq pipeline | https://zenodo.org/record/156534 | https://github.com/ENCODE-DCC/atac-seq-pipeline |
| deeptools (ver 3.5.1) | https://doi.org/10.1093/nar/gkw257 | https://deeptools.readthedocs.io/en/develop/ |
| Homer (ver 4.11) | Benner Lab | http://homer.ucsd.edu/homer/ngs/ |
